# Recovery of microbial ecophysiology and carbon accrual functions in peatlands under restoration

**DOI:** 10.1101/2025.08.01.668219

**Authors:** William Pallier, Yiling Wang, James C. Kosmopoulos, Rebekka R.E. Artz, Claire McNamee, Ezra Kitson, Nicholle Bell, Andreas Richter, Roxane Andersen, Thomas C. Parker, Ashish A. Malik

**Affiliations:** School of Biological Sciences, University of Aberdeen, Aberdeen, UK; School of GeoSciences, University of Edinburgh, Edinburgh, UK; Department of Bacteriology, University of Wisconsin-Madison, Madison, WI, USA; Microbiology Doctoral Training Program, University of Wisconsin-Madison, Madison, WI, USA; Ecological Sciences, The James Hutton Institute, Aberdeen, UK; School of Chemistry, University of Edinburgh, Edinburgh, UK; Centre for Microbiology and Environmental Systems Science, University of Vienna, Vienna, Austria; Environmental Research Institute, University of Highlands and Islands, Thurso, UK

## Abstract

Peatlands are water-logged ecosystems that limit microbial decomposition making them effective carbon sinks. However, drainage or erosion removes these constraints on decomposition, switching them to carbon sources. Restoration aims to reverse these trends. Microbial ecophysiology influences carbon fluxes but how it responds to peatland degradation and restoration is poorly understood. Here we used metagenomics to study microbial functions and quantified growth rates using isotope labelling across seven sites in Britain, each with restored, degraded, and near-natural peatlands. We found that growth rates in restored treatments were comparable to the near-natural, but were significantly higher in degraded. This growth rate reduction in restored peatlands was dependent on the scale of degradation and the length of restoration, and was underpinned by a shift towards energetically less favourable metabolic pathways such as anaerobic respiration, fermentation, and carbon fixation. A peatland ecosystem health index estimated based on measurements of peat moisture, oxygen, pH, organic matter chemistry, and moss cover, explained a significant amount of variation in microbial ecophysiology across the gradient. We demonstrate that microbial ecophysiology changes with peatland ecosystem health in a predictable manner. This knowledge can inform restoration targets and monitoring of recovery to maximise the return of carbon accrual functions of peatlands.

## Main

Peatlands cover only ∼3% of the land area on Earth yet store ∼21% of the global total soil carbon stock (Yu et al., 2010; Köchy et al., 2015; Poulter et al., 2021). However, around 10% of peatlands globally have been drained, land cover-converted, or mined (Turetsky et al., 2015; Joosten et al., 2016; Leifeld and Menichetti, 2018; Xu et al., 2018). This creates aerobic conditions that lead to a faster mineralisation of organic matter, increasing emissions of greenhouse gases such as carbon dioxide (CO_2_) and shifting peatlands from being net carbon sinks to carbon sources (Tiemeyer et al., 2016; Leifeld et al., 2019; Evans et al., 2021). Each year drained peatlands emit ∼2 Gt of carbon dioxide which, according to some estimates, are in the range of 5% of anthropogenic greenhouse gas emissions (Joosten et al., 2016; Huang et al., 2021). Therefore, peatland protection and restoration efforts have become an area of focus in national strategies to meeting climate mitigation targets (Leifeld & Menichetti, 2018).

The amount of carbon stored in natural peatlands is the balance of carbon inputs from photosynthesis and carbon losses through respiration, a large proportion of which is driven by microbial decomposition. The abundance, diversity, physiology, and interactions of these microbes play key roles in controlling the dynamics of carbon and nutrients in soils (Schimel and Schaeffer, 2012; Sokol et al., 2022) including peatlands (Geay et al., 2024; Gios et al., 2024). Their growth and activity, or lack of, governs the fate of belowground carbon inputs (Sokol et al., 2022). Therefore, better management of peatland restoration requires a mechanistic understanding of microbial ecophysiology and biogeochemistry. However, most studies of peatland microbes in restoration ecology have focussed on studying taxonomic changes (Andersen et al., 2012; Kitson and Bell, 2020) while understanding remains poor of the exact microbial physiological mechanisms involved in organic matter decomposition studied through the lens of microbial genes and genomes.

In peatlands, complex feedback mechanisms between ecology, biogeochemistry and hydrology regulate carbon cycling (Holden et al., 2004; Waddington et al., 2015). Hydrology influences the vegetation community, belowground carbon inputs and rates of decomposition (Zhong et al., 2020). In a simplistic view, a high water table leads to waterlogged conditions and a reduction in oxygen permeability in peat soils; drainage lowers the water table which decreases the soil moisture content thereby increasing the number of air-filled pores and allowing oxygen to permeate deeper (Silins and Rothwell, 2011; McCarter et al., 2020). Lack of oxygen curtails microbial aerobic respiration, a high energy yielding pathway. As a consequence, microbes in peat soils must use alternative terminal electron acceptors with the most common anaerobic respiration reactions in the following order of energy yield: denitrification, manganese reduction, iron reduction, sulphate reduction and methanogenesis (Wang et al., 2017; Zhang and Furman, 2021). Additionally, fermentation is used to generate energy using monosaccharides under anoxic conditions. Both anaerobic respiration and fermentation are thermodynamically less favourable than aerobic respiration which results in slower microbial growth rates and activity and consequently a reduction in decomposition rates (Laiho, 2006). Metagenomic studies have revealed that microbes in peatlands have a mix of aerobic and anaerobic metabolism, particularly in the highly dynamic upper layers (Lipson et al., 2013; Lin et al., 2014). The lack of oxygen is considered the main constraint on organic matter decomposition in peatlands, but we do not know the scale of shift in microbial ecophysiology, growth rates, rates of decomposition, and carbon flux with water table variations.

Peatland hydrology and vegetation are also tightly coupled. In the temperate oligotrophic peatlands studied here, *Sphagnum* dominates and contributes to maintain wet conditions, while vascular vegetation become more dominant in drier conditions; this shift affects the rooting depths thereby introducing oxygen and more labile organic matter deeper into the peat profile (Gunnarsson, 2005; Strack and Waddington, 2007; Potvin et al., 2015; Rajakaruna et al., 2024). The shift in vegetation also leads to varying organic matter inputs alongside its composition which in turn affects decomposition by microorganisms (Zeh et al., 2020; Wang et al., 2021). Organic matter with high soluble carbon compounds and low lignin or aromatic compounds is decomposed faster; in peatlands, sedge-derived organic matter decomposes relatively faster compared to mosses while woody plant litter is intermediate (Laiho, 2006; Andersen et al., 2013; Bengtsson et al., 2018). Therefore, in addition to oxygen availability, the vegetation type and peat organic matter composition influences microbial ecophysiology in terms of capabilities to degrade different substrate types which influences rates of decomposition and the ability of peatlands to store carbon.

Given the increased pressures on peatlands associated with rapidly changing climate (Turetsky et al., 2011, Turetsky et al., 2015; Swindles et al., 2019; Wilkinson et al., 2023) and the increased pace and scale of restoration interventions (Andersen et al., 2017; Loisel et al., 2021; Rowland et al., 2021), there is an urgent need to better understand how microbial ecophysiology shifts following peatland restoration and determine how these changes in microbial ecophysiology can impact emergent functions and ecosystem processes. Knowledge of these processes and how they are influenced by shifts in environmental conditions that accompany peatland degradation and restoration is required to both predict and manage carbon accrual in peatlands of varying ecosystem health (Ritson et al., 2021). Such knowledge allows fine tuning of models with better functional understanding that is crucial to determine the target sites for restoration to achieve maximum benefits and to monitor progress of restoration in recovery of microbial and consequent ecosystem functions. This will improve the success of restoration efforts in securing carbon stocks and returning the carbon sink functions of peatland ecosystems.

The overall aim of this study was to assess the shifts in microbial ecophysiology in restored peatlands relative to the degraded and reference near-natural contrasts, in order to monitor the efficiency of restoration in recovering microbial carbon cycling functions. To achieve this, we used an observational landscape-scale study of seven peatland sites across Britain, each containing locally adjacent restored, degraded and near-natural (best available target state) areas. We quantified key microbial carbon cycling traits in the top peat layer (0-10 cm) using metagenomics, measured genome-level pathways as potential for different metabolic and substrate degradation functions of bacteria and archaea and quantified the consequences for community-scale microbial growth rates using isotope labelling. We hypothesised that the decline in oxygen levels in restored compared to degraded peatlands imposes constraints on microbial growth and reduces their carbon cycling rates. We also generated an ecosystem heath index using measured environmental variables and vegetation cover to ascertain whether shifts in microbial ecophysiology can be explained using these simple measures as a proxy. By combining these, we provide a better understanding of the constraints on microbial processes and decomposition in peatlands, the impacts of peatland degradation and degree of success in restoring peatland microbial and ecosystem functioning.

## Results

### Restoration leads to a partial recovery of environmental characteristics

The environmental characteristics of the surface peat samples from near-natural (hereafter natural for simplicity), degraded, and restored peatlands from the seven sites (Figure 1a, Table S1) demonstrated higher similarity by site, explaining 39% of variance, than by treatment categories of ecosystem health which explained 15% of variance. The interaction between site and treatment was also significant, explaining 20% of variance, suggesting that treatment effects may vary by site (Figure 1b). This variation across the samples along the first two axes of Principal Component Analysis (PCA) was represented by six factors (Figure 1b) in the following order: dissolved oxygen concentration (19.8%), C:N ratio (19.1%), organic matter chemical composition as derived from the first PCA axis of Fourier Transform Infrared or FTIR spectroscopy (18.1%), moisture content (17%), moss cover (13.5%) and pH (12.4%). Other measured environmental variables namely bulk density, carbon and nitrogen concentration, carbon stock, vascular plant cover, and electrical conductivity were excluded from this analysis due to collinearity with one or more of the six key factors.

**Figure 1.**
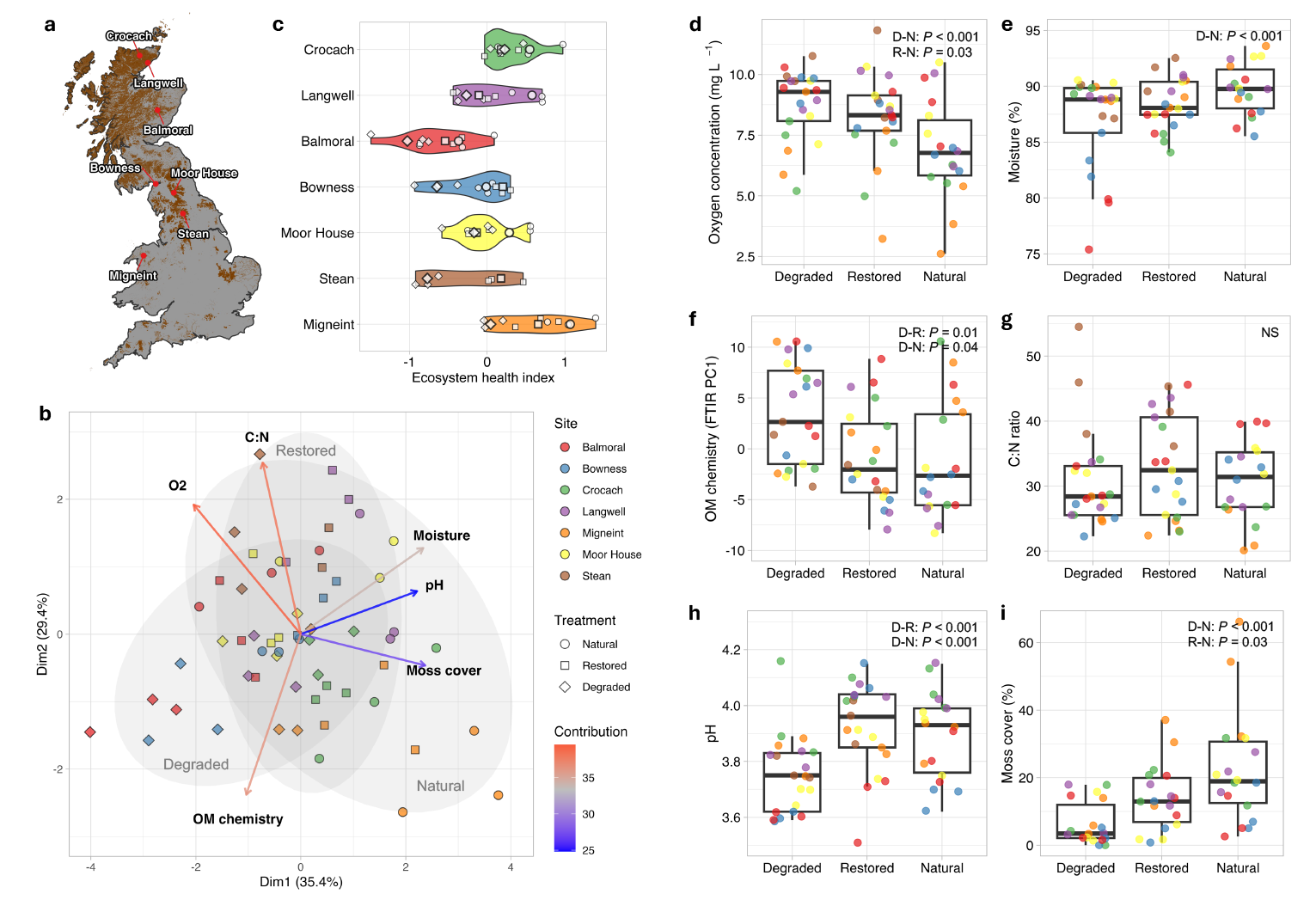
Study sites and ecosystem properties. a) Geographical distribution of the study sites over a map of Britain with the extent of peat cover shaded in brown. b) Principal component analysis (PCA) plot of the environmental variables across all samples. The vectors represent significant environmental variables, their direction represents an increasing trend, and their colour represents contribution to the total variation in the first two principal components. Shaded areas cover the sample variation across treatments with 90% confidence intervals. c) Site-wise distribution of the ecosystem health index calculated for each sample with the larger points representing the mean values across the treatments. The index spans from negative to positive values representing increasing ecosystem health. d-i) Environmental variables for all samples shown across the treatments of degraded, restored and natural peatlands for d) oxygen concentration, e) moisture content, f) organic matter chemical composition represented by FTIR PC1, g) C:N ratio of the organic matter, h) peat pH, and i) proportion of moss cover in the aboveground vegetation. Only significant pairwise differences in environmental variables across the three treatments are highlighted with *P* values calculated using mixed effects models with site as a random effect (D: degraded, R: restored, and N: natural, NS: not significant). Thick horizontal lines in the boxplots show median, boxes show interquartile ranges and vertical whiskers extend 1.5 times the inter quartile range.

Despite strong site-level differences, we could clearly identify broad patterns in the environmental characteristics of our predefined ecosystem health categories of natural, restored, and degraded peatlands. Peatland degradation clearly results in substantial system-level changes; the water table was lowered (Figure S1) resulting in less wet, more oxygenated surface peat (Figure 1d-e, S2). These changes cause peat compaction as observed by the higher bulk density in degraded peatlands compared to restored and natural treatments (Figure S3). Carbon and nitrogen concentration in peat across treatments did not vary significantly but topsoil (0-10cm) carbon stock was higher in degraded compared to restored and natural treatments (Figure S3) due to higher bulk density. Degraded peatlands generally store less carbon, but such an assessment of total carbon stocks requires sampling the entire peat profile.

The chemical composition of peat organic matter measured using FTIR spectroscopy (PCA first axis which explained 50.7% variation) varied across our treatments of ecosystem health (Figure 1f); organic matter composition in degraded peatlands varied from natural and restored peatlands, whereas it was similar in natural and restored peatlands. The C:N ratio of peat organic matter was not statistically different across treatments (Figure 1g). The overall trends in organic matter composition and C:N ratio hints at higher decomposition rates and turnover of organic matter in degraded peatlands (Kalbitz and Geyer, 2002; Krüger et al., 2015; Leifeld et al., 2020). Soils in degraded peatlands were more acidic with an average pH of 3.75 compared to 3.89 in near-natural peatlands and 3.92 in restored peatlands (Figure 1h). Acidification on degradation has been widely reported in peatlands due to increased accumulation of degradation products (Laiho, 2006; Xue et al., 2023), but this can be context dependent. Across our treatments of ecosystem health, there was also a shift in the vegetation composition from moss dominated to vascular plants (Figure 1i). The observed environmental variables in our restored peatlands with raised water table hinted at a good recovery from degradation, but they still differed from the natural reference. These changes in environmental characteristics with varying water table have been well documented in numerous wetland studies (Elliott et al., 2015; Urbanová and Bárta, 2016; Klimkowska et al., 2019; Yang et al., 2025).

### An ecosystem health index captures variation across sites and treatments

To tease apart the site level differences, we used the six key environmental parameters explaining variation across samples to estimate an ecosystem health index (Figure 1c). The index was higher when oxygen concentration and organic matter chemical diversity was lower, and moisture, C:N ratio, pH and moss cover was higher all representing better peatland ecosystem state. The index differed significantly between degraded, restored and natural treatments with lower, intermediate and higher mean values, respectively (Figure S4). When it was tested across our ecosystem health categories at individual sites there were significant differences in four of the seven sites although the general trend was similar across all sites (Figure S5) which highlights that restoration improved the ecosystem state. Much of the variation in the index across samples was likely driven by site-level differences in peat type, legacy pressures and climate. An idealised state of ecosystem health might be unachievable in the majority of peatlands because of their inherent location in the climatic, legacy management, and hydrological landscape but shifts in the index could be used to monitor improvements in ecosystem state while comparing damaged, restored and natural areas within a peatland site. Across our sampled sites, Migneint, Crocach and Langwell demonstrated higher ecosystem health, followed by Moor House, Bowness and Stean, and Balmoral the lowest. Balmoral and Stean peatlands are highly eroded whereas Crocach and Langwell, the sites in northern Scotland, are among the best reference peatlands which suggests that the ecosystem health index is capturing meaningful variation. The age of restoration also varied across sites, it was 3 years for Balmoral, 6 years for Langwell and 9-12 years for the other sites. Overall, we show that restoration produces substantial changes in the environmental characteristics of surface peat soils but results in peatlands where the legacy of drainage can persist. Such an index has been used in agricultural context to evaluate soil health (Amacher et al., 2007; Lehmann et al., 2020; Li et al., 2021) or land use intensity (Blüthgen et al., 2012) but has never been applied in peatlands at the field scale (Worrall et al., 2025). This environmental gradient across the ecosystem health categories allowed us to study the effect of restoration on the functional recovery of soil microorganisms. In addition to assessing the differences across the ecosystem health categories, we also used the ecosystem health index to highlight site level differences in microbial functions and make generalizable predictions.

### Reimposing of constraints on microbial growth in restored peatlands

To test our key hypothesis that the reduction in oxygen in restored compared to degraded peatlands imposes constraints on the growth, i.e., cell division, of microorganisms and therefore slows their carbon cycling rates, we measured community-level growth rates using ^18^O incorporation from isotopically labelled water into microbial total DNA in lab incubations at a standard temperature and field moisture conditions. Potential growth rates were significantly higher in degraded peatlands when compared to natural and restored (Figure 2a). Rates in restored treatments did not differ significantly from the natural peatlands suggesting reimposing of constraints on microbial growth potentially reducing rates of decomposition and CO_2_ flux when compared to degraded peatlands.

**Figure 2.**
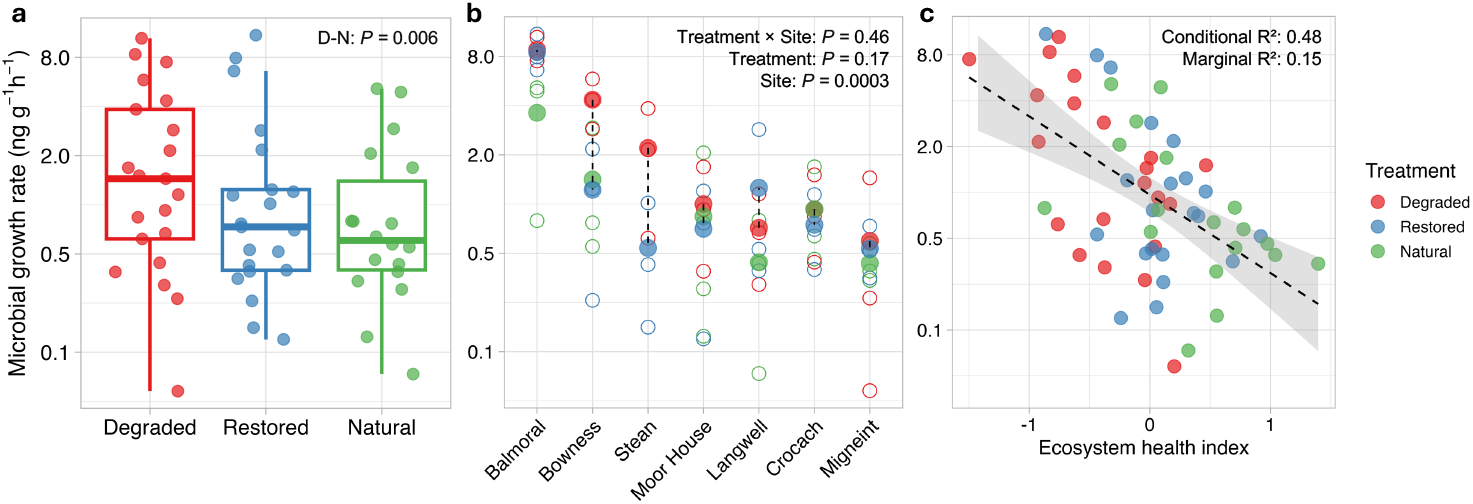
Microbial community-level growth rates. a) Community-aggregated potential growth rates measured using ^18^O incorporation into DNA from isotopically labelled water across the three treatments in all sites. Only significant pairwise differences in growth rates across the three treatments are highlighted with *P* values calculated using mixed effects models with site as a random effect (D: degraded, R: restored, and N: natural). Thick horizontal lines in the boxplots show median, boxes show interquartile ranges and vertical whiskers extend 1.5 times the inter quartile range. Growth rates are plotted on a log_2_ scaled axis. b) Growth rates presented across the seven sites; all samples are shown as empty points whereas the filled points denote the mean values for each treatment. The dashed vertical line connects the mean of restored with degraded treatment to highlight the change in growth rates. *P* values calculated using a mixed effects model for site, treatment, and their interaction are shown. c) Growth rates regressed against ecosystem heath index for all samples. The dashed line fits a linear regression model to the data with the shaded area showing a 95% confidence interval; the marginal and conditional R^²^ from a linear mixedeffects model are shown representing the variation in microbial growth due to ecosystem health index alone and due to both ecosystem health and site-level variation, respectively.

To test how microbial potential growth rates varied across the seven sites in addition to the ecosystem health treatments, we performed linear mixed effects modelling and observed that site was the most significant factor explaining 46% of the variation in growth rates. Balmoral, Bowness and Stean sites which are significantly degraded relative to the other sites demonstrated the highest growth rates followed by Moor House, Langwell, Crocach, and Migneint (Figure 2b). In some sites, we observed a trend that the degraded treatment had higher growth rate relative to the restored, and the natural reference had the lowest. The reduction in growth rates from degraded to restored treatments suggests the mitigation effect of restoration and this was very clear in Bowness and Stean where the damaged sites are in a really poor condition and restoration has been ongoing for roughly a decade. Such a reduction in growth rate was not seen in Balmoral, also in a really poor condition but the restoration age was only 3 years. Other sites with better ecosystem health already have lower growth rates and restoration did not bring about major changes. This is further emphasised by the observed significant negative regression of growth rate with ecosystem health index across all sites (Figure 2c). The ecosystem health index alone explained 15% of the variation in microbial growth rates; when site is considered as a random factor nearly half the variation is growth rate is explained by the ecosystem health index. When tested within each site, a similar negative trend was observed but this was weaker in the sites with better ecosystem health (Figure S6).

When we tested the influence of the six key environmental factors on the variation in growth rates using a combined multiple linear regression model (Figure S7), moisture was identified as the main co-variate (*P*<0.001) followed by C:N ratio (*P*=0.005) and pH (*P*=0.02). Oxygen which was hypothesised as the main driver of microbial growth rate did not covary significantly with growth rate in the combined model, but when tested individually it varied significantly (*P*=0.02). Overall, these phenotypic measurements of microbial ecophysiology highlights the concurrent scale of potential change in microbial and ecosystem processes accompanying ecosystem health degradation and restoration in peatlands. Measurements of changes in microbial growth rates in peat with changing water table are rare (Bergman et al., 2000; Bell et al., 2023) and here we provide estimates of differences across peatlands of varying ecosystem health thereby giving insights into the recovery of microbial ecophysiology on restoration across multiple sites.

### Variation in prokaryotic communities in peatlands explained by its ecosystem health

To probe microbial recovery on restoration, we used shot gun metagenomics in surface peat soils comparing natural reference with restored and degraded treatments of ecosystem health. Using assembly and binning approaches we retrieved 935 Metagenome-Assembled Genomes (MAGs) that were estimated to be of medium or high quality. We retrieved 862 Bacterial MAGs from 21 phyla and 73 Archaeal MAGs from 5 phyla. We first clustered these 935 MAGs to generate 459 species-level representative MAGs for taxonomic analyses. Using principal coordinate analysis (PCoA), we found that the prokaryotic community composition was driven more by site (R^2^=0.36, *P*<0.05) than ecosystem health categories (R^2^=0.04, *P*<0.05), although the interaction of site and treatment was also significant (R^2^=0.43, *P*<0.05) highlighting that the effect of ecosystem health on prokaryotic community composition varied by site (Figure 3a). PCo1 which explained 19.11% of the variation in prokaryotic community composition was associated with the ecosystem health index as demonstrated using a regression of the two across all sites (Figure 3b; R^2^=0.3, *P*<0.001). A similar negative trend was seen in five of the seven sites (Figure S8). Within each site, the effect of ecosystem health can also be observed; diversity differences across the ecosystem health treatments at individual sites were significant in all but one site (Figure 3c). While microbial communities from the restored treatments appeared distinct from the degraded and natural across most sites, sites that were more degraded (Balmoral, Bowness and Stean) were compositionally similar to each other and different from sites with better ecosystem health (Langwell and Mignient) with one site (Moor House) demonstrating high variability. Overall, our results suggest that the prokaryotic community assembly in peatlands can be driven by its ecosystem health in a somewhat predictable manner.

**Figure 3.**
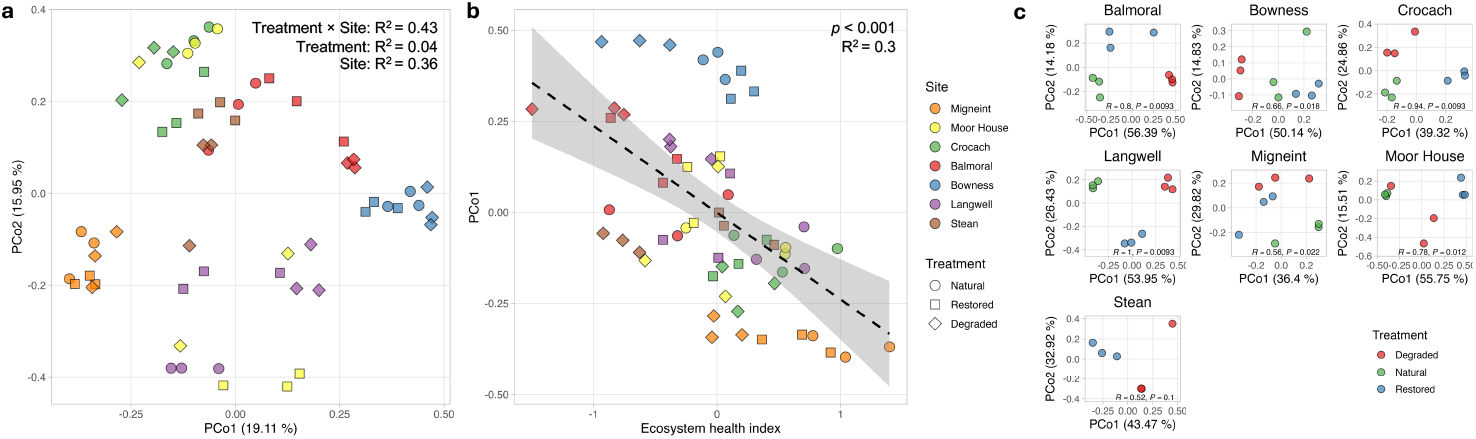
Variation in microbial community assembly: a) Bray Curtis distance-based Principal Coordinate Analysis (PCoA) plot of microbial community composition based on variation of MAG abundance across all samples. Colour coding as in figure 3b. R^²^ from PERMANOVA analysis are provided for site, treatment, and their interaction. b) PCo1 from the PCoA analysis regressed against ecosystem heath index for all samples. The dashed line fits a linear regression model to the data with the shaded area showing a 95% confidence interval; adjusted R^²^ and *P* value from a linear mixed-effects model are shown representing the variation in PCo1 with the ecosystem health index with site as a random factor. c) PCoA plot of microbial community composition when analysed separately for each site showing variation of MAG abundance across the treatments. R statistics and *P*-values derived from analysis of similarity (ANOSIM) report the degree of separation for communities from different treatments.

### Shifts in energy metabolism and substrate degradation along peatland ecosystem health

We measured assembled contigs-derived community functional gene abundances (normalised by total predicted protein-coding genes) for metabolic pathways of interest and decomposition genes for key substrates. We used gene annotations using Kyoto Encyclopaedia of Genes and Genomes (KEGG) which encompasses pathways of both aerobic and anaerobic metabolism. We also used the Carbohydrate Active Enzymes (CAZy) database which contains gene information for enzymes involved in degradation of carbohydrates and other substrates. We first focussed on the key pathways for energy metabolism that are most relevant to peatland soil microorganisms, namely fermentation, carbon fixation, and metabolism of nitrogen, sulphur and methane (Zhang and Furman, 2021). We observed that microbial metabolism was less anaerobic in degraded and restored peatlands compared to the reference natural, an observation widely reported in numerous studies from peatlands (Lipson et al., 2013; Lin et al., 2014; Zhang and Furman, 2021). For anaerobic processes lower down the redox ladder, such as methanogenesis, the abundance of methane metabolism genes was significantly higher in the restored and reference natural compared to the degraded treatment (Figure 4e). For the other key metabolic pathways of interest, while there was no significant variation across categories of ecosystem health (Figure 4a-d), the community-level gene abundance significantly increased along the ecosystem health index (Figure 4f-j) reinforcing the relevance of these pathways in intact peatlands. While gene abundances for all five metabolic pathways of interest increased significantly along the ecosystem health index, fermentation, carbon fixation, and methane metabolism pathways were the better predicted using the index than nitrogen and sulphur metabolism highlighting their significance in the ecosystem functioning of the peatlands under study. When we tested the variance of gene abundances for all five metabolic pathways along the ecosystem health index for each site individually, we observed that the slope was larger in the degraded sites (Figure S9). A negative regression of gene abundances for the metabolic pathways of significance in our sites (carbon fixation, methane metabolism, and fermentation) against growth rate was also observed (Figure S10). Our results demonstrate that a shift away from these signature peatland metabolic processes to energetically more favourable aerobic respiration in peatlands with lower ecosystem health underpins the observed increase in microbial growth rates and therefore higher potential rates of decomposition.

**Figure 4.**
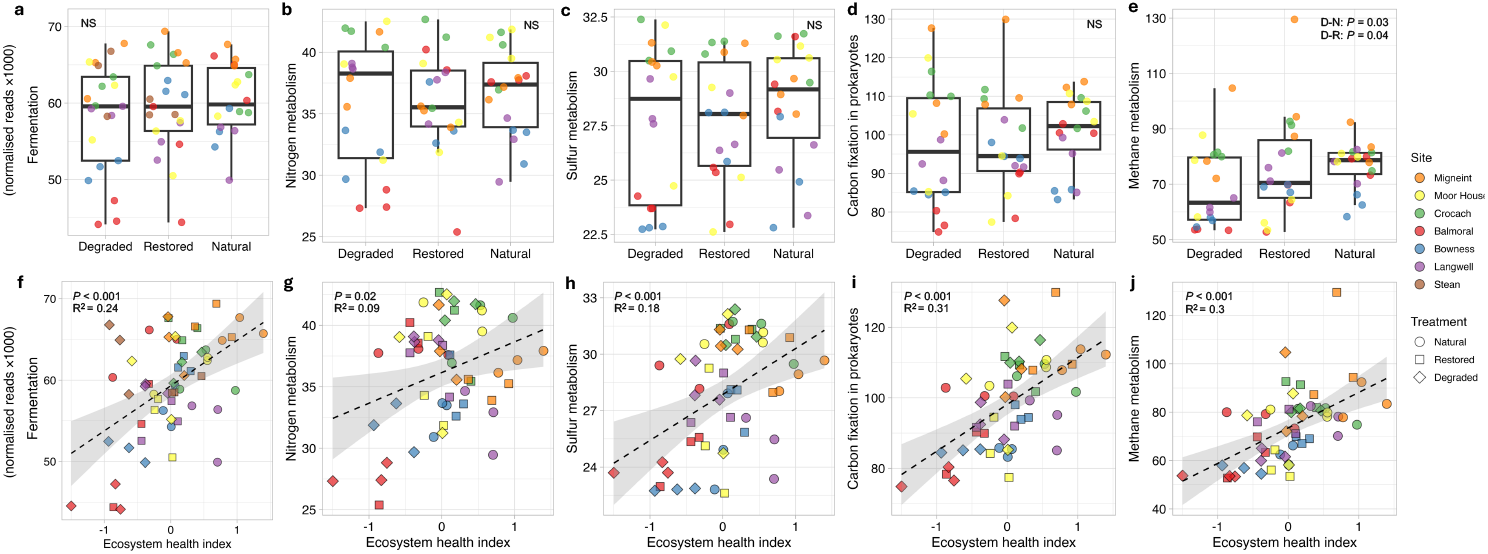
Microbial community-level metabolic functions: Variation across treatments of sum abundance of genes as normalised reads (×1000) belonging to key functional categories of metabolic pathways: a) fermentation, b) nitrogen metabolism, c) sulphur metabolism, d) carbon fixation in prokaryotes, and e) methane metabolism. Only significant pairwise differences in gene abundances across the three treatments are highlighted with *P* values calculated using mixed effects models with site as a random effect (D: degraded, R: restored, and N: natural, NS: not significant). Thick horizontal lines show median; boxes show interquartile ranges and vertical whiskers extend 1.5 times the inter quartile range. Gene abundances regressed against ecosystem heath index for all samples for f) fermentation, g) nitrogen metabolism, h) sulphur metabolism, i) carbon fixation in prokaryotes, and j) methane metabolism. The dashed line fits a linear regression model to the data with the shaded area showing a 95% confidence interval; adjusted R^²^ and *P* value from a linear mixed-effects model are shown. The Stean site was excluded from this analysis due to very high level of variation causing the samples to appear as significant outliers.

Microorganisms in the natural reference, degraded, and restored peatlands varied significantly in their ability to decompose carbon substrates. Degraded and restored peatlands in comparison to the natural have increased abundance of genes associated with depolymerisation of more recalcitrant aromatic compounds (Figure 5d). In contrast, natural reference have greater abundance of genes to break down simpler carbon compounds such as oligosaccharides when compared to degraded and restored peatlands (Figure 5c). Degraded and restored peatlands also have greater abundance of genes involved in depolymerising microbial cell walls (Figure 5e) which implies an increased recycling of microbial necromass in these systems which can be clearly linked to the observed increased microbial growth rates and potential carbon turnover in these systems. We also linked gene abundance to the ecosystem health index and observed that oligosaccharides and cellulose depolymerising genes increased with higher ecosystem health (Figure 5g-h), whereas aromatic substrate decomposition genes decreased with higher ecosystem health (Figure 5i). Genes for lignin decomposition did not show any discernible trend (Figure 5a,f). These trends varied when regressed against ecosystem health index separately for each site (Figure S11) suggesting that differences across sites mostly drove the variation in gene abundance for substrate degradation along our gradient of ecosystem health.

**Figure 5.**
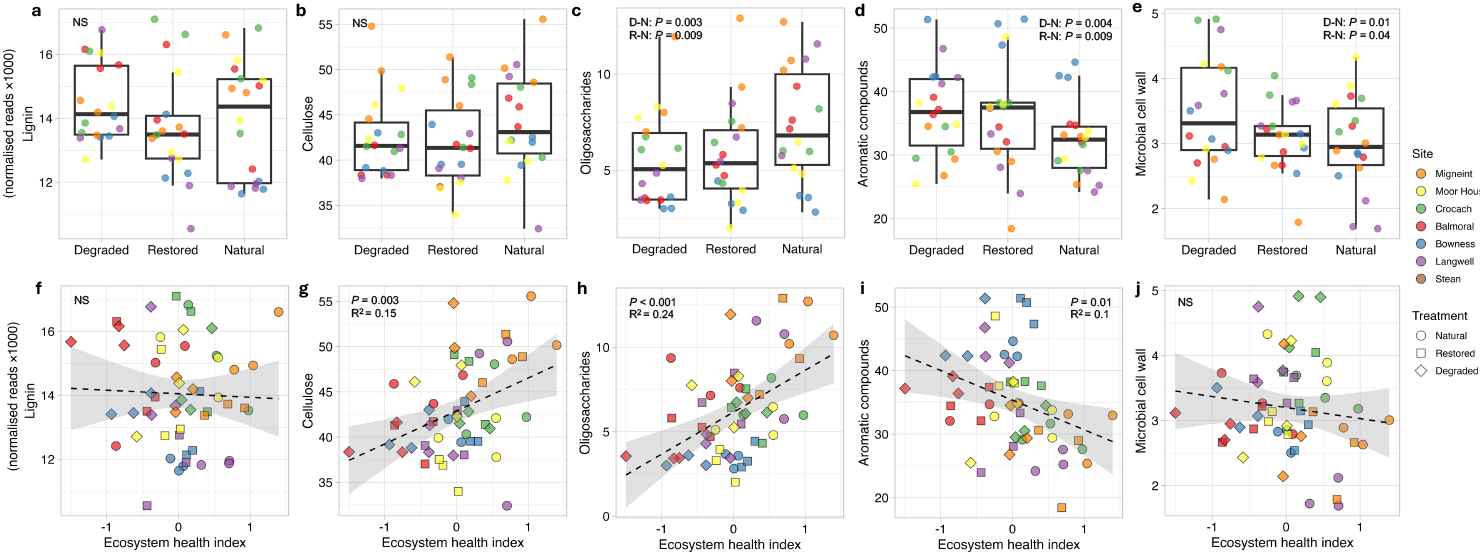
Microbial community-level substrate decomposition functions: Variation across treatments of sum gene abundance as normalised reads (×1000) for decomposition of the following substrates: a) lignin, b) cellulose, c) oligosaccharides, d) aromatic compounds, and e) microbial cell wall. Only significant pairwise differences in gene abundances across the three treatments are highlighted with *P* values calculated using mixed effects models with site as a random effect (D: degraded, R: restored, and N: natural, NS: not significant). Thick horizontal lines show median; boxes show interquartile ranges and vertical whiskers extend 1.5 times the inter quartile range. Gene abundances regressed against ecosystem heath index for all samples the following substrates: f) lignin, cellulose, h) oligosaccharides, i) aromatic compounds, and j) microbial cell wall. The dashed line fits a linear regression model to the data with the shaded area showing a 95% confidence interval; adjusted R^²^ and *P* value from a linear mixed-effects model are shown. The Stean site was excluded from this analysis due to very high level of variation causing the samples to appear as significant outliers.

The increase in community-level abundance of genes for decomposition of more complex substrates in degraded and, to some extent, in restored peatlands can be linked to the observed changes in the peat organic matter composition (Figure 1f). FTIR derived abundances of carbohydrates, and aliphatic and aromatic compounds were significantly higher in degraded peatlands in comparison to the natural reference and restored treatments. These changes in organic matter following peatland degradation likely occurs through increased microbial decomposition leading to the accumulation of aromatic compounds and the loss of low molecular weight compounds, and through shifts in carbon inputs from changes in the vegetation composition. Such changes in the organic matter chemical composition in response to water table changes have been widely reported (Artz et al., 2008; Krüger et al., 2015; Negassa et al., 2021) but here we link them to substrate decomposition capabilities of the microbial communities. We observed that the microorganisms in restored peatlands still have the genetic capability to degrade complex carbon compounds, which means that if the immediate constraints on microbial metabolism and growth (through lower oxygen concentrations due to higher water table) are removed, increased decomposition will likely resume. This is plausible due to the dynamic nature of the water table. Also, increased drought occurrences (Swindles et al., 2019) are likely to create phases when constraints on microbial metabolism and growth will be reduced causing increased decomposition of complex substrates. This hints at a lower resistance of restored peatlands compared to the natural reference to becoming carbon sources when water table is transiently lowered.

### Genomic metabolic capabilities under different ecosystem health

We then probed for genome-level capabilities as proxies of microbial functional potential across the peatland ecosystem health treatments. By mapping the absolute metagenomic read counts to species-representative bacterial and archaeal MAGs followed by differential abundance analysis and hierarchical clustering, we identified populations that were enriched in each of the three treatments within each site. We found that 39% of the total MAGs were enriched in the natural treatment which was higher than the 34% in damaged and 27% in restored. We then explored these enriched MAGs found in the three treatments for presence or absence of our five metabolic pathways of interest: fermentation, carbon fixation, and metabolism of nitrogen, sulphur and methane (Figure 6).

**Figure 6.**
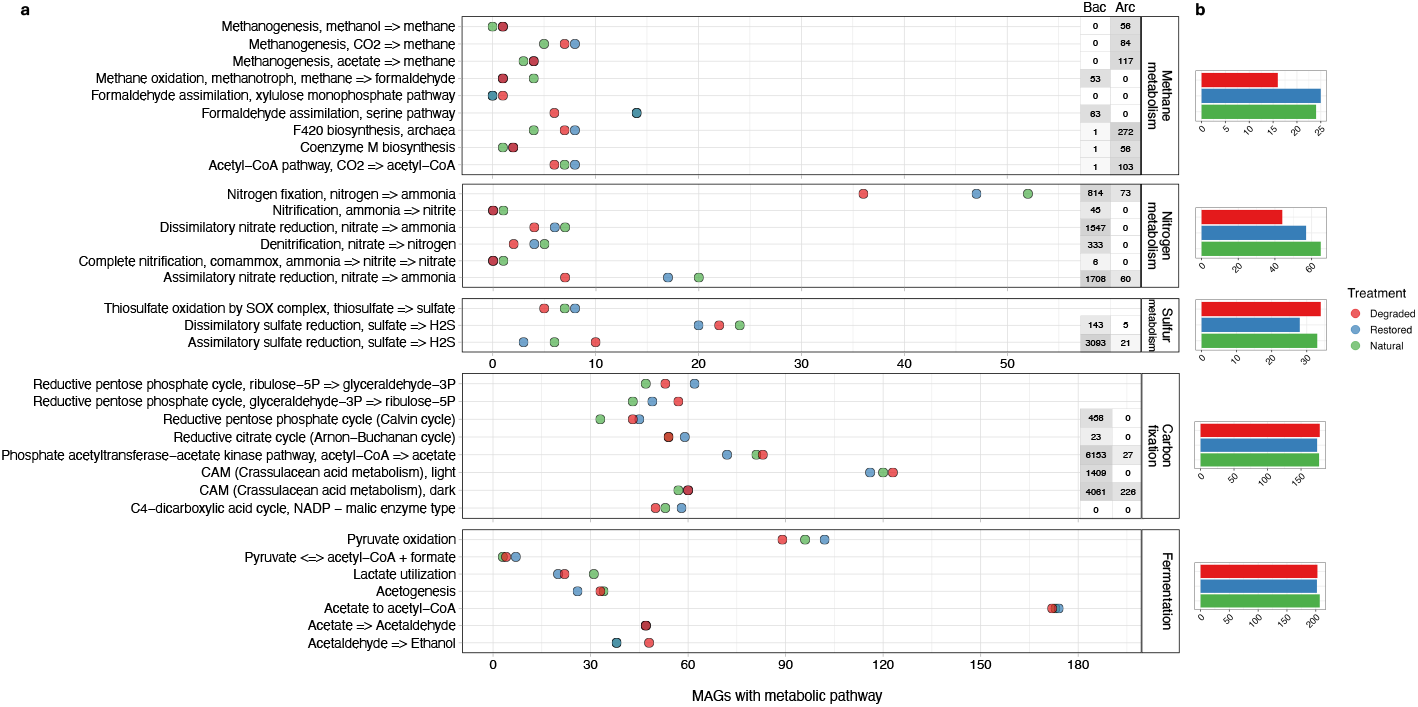
Microbial population-level metabolic potential: a) Number of unique MAGs that were differentially abundant across the treatments with presence of metabolic modules within the five metabolic pathways of interest. The numbers at the side for each module represents the total number of bacteria (Bac) and archaea (Arc) in the KEGG database with a complete gene set for the module as an indication of pathway presence across the tree of life. Some points overlap which can be seen by a darker shade. b) Sum number of unique MAGs across the treatments with the presence of at least one metabolic module

We observed a higher number of enriched bacteria and archaea with methane metabolism capabilities in the restored and natural compared to degraded peatlands. Specifically, higher number of enriched methanotrophic bacterial MAGs with capability for assimilation of methane-derived formaldehyde through the serine pathway (Averesch and Kracke, 2018) were identified in restored and natural treatments compared to the degraded (Figure 6a). Enriched archaeal MAGs were found that coded for functions such as methanogenesis from CO_2_, F_420_ biosynthesis which is involved in key steps of methanogenesis (Grinter and Greening, 2021), and acetyl-CoA pathway which converts CO_2_ to acetyl-CoA which can then be used in methanogenesis with hydrogen or formate (Martin, 2020), but their numbers did not vary across the treatments highlighting similar functional capabilities. Higher number of enriched bacterial and archaeal MAGs in natural followed by restored followed by degraded had functional capabilities for different nitrogen metabolism pathways. This trend was most noticeable for nitrogen fixation (conversion of nitrogen to ammonia) which was generally very widespread across all treatments; biological nitrogen fixation rates have been reported to be higher than the rates of nitrogen deposition in northern peatlands making it a significant pathway of nitrogen input to the system (Yin et al., 2022). Similarly, assimilatory nitrate reduction (conversion of nitrate to ammonia) was also present in more enriched bacterial and archaeal MAGs in natural and restored compared to degraded peatlands. There was no clear trend in sulphur metabolism capabilities; with dissimilatory sulphate reduction (conversion of sulphate to hydrogen sulphide) being more widespread than other sulphur metabolism pathways (Pester et al., 2012).

Carbon fixation capabilities were equally widespread across all three treatments (Figure 6b) although reductive pentose phosphate cycle was more prevalent in enriched bacterial MAGs in degraded and restored compared to natural peatlands. Crassulacean acid metabolism which represents plant-like fixation of carbon dioxide through phosphoenolpyruvate carboxylase although an incomplete pathway in bacteria and phosphate acetyltransferase-acetate kinase pathway which coverts acetyl-CoA to acetate were the most common carbon fixation pathways across treatments. Carbon fixation by photoautotrophs, chemoautotrophs as well as mixotrophs can be quite significant in peatlands contributing substantially to the ecosystem carbon balance (Hamard et al., 2025) but the metabolic pathways involved have been understudied. Fermentation was widespread across treatments with acetate to acetyl-CoA conversion and pyruvate oxidation being the most common metabolic pathways. Collectively, methane, nitrogen and sulphur metabolism capabilities were present in fewer enriched prokaryotes than carbon fixation and fermentation pathways (Figure 6b), reflecting the greater phylogenetic conservation of the former pathways compared to the latter. Overall, our genome-level analysis of enriched bacterial and archaeal populations from the three peatland treatments sheds light on their unique metabolic capabilities and opens avenues for further research on how peatland microbes driving key biogeochemical cycles shift in response to land management change and climate extremes with implications for ecosystems functioning.

## Discussion

We observed that across a gradient of temperate oligotrophic peatland sites varying in their apparent ecosystem health, microbial genomic and phenotypic traits were predictable using an ecosystem health index based on measurements of moisture, dissolved oxygen, pH, organic matter C:N ratio and chemical composition, and moss cover. Across the seven sites that were studied each with the three categories of ecosystem health, distribution of microbial traits was better explained by the ecosystem health index that by the ecosystem health categories. The index enabled this because of incorporation of site level differences due to peat type, climate, and legacy land use pressures which makes it generalisable to predict patterns of microbial functional changes following peatland degradation and restoration. Such a health rating has been successfully tested for agricultural soils to infer the genomic traits of bioindicators which were evaluated for their environment-wide associations with differing management practices (Wilhelm et al., 2023). In peatlands, microbial community structure and functions have been predicted using information on *Sphagnum* phylogeny, anatomical and morphological traits, and metabolites, with metabolites being more influential than other factors (Sytiuk et al., 2022). While microbial genomics tools are not ideal candidates to monitor ecosystem health due to the complex nature of the generated data and high costs, our results highlight the applicability of the ecosystem health index in predicting microbial functions. This can inform restoration targets depending on the scale of peatland degradation and monitoring of recovery under restoration so that we can maximise the return of carbon accrual functions in peatlands. The index measured using simple environmental variables needs further validation to ascertain whether it provides information on key ecological, diversity and biogeochemical cycling attributes of a healthy peatland ecosystem.

Microbial community-level growth rates were much lower in the healthier sites and treatments, along with higher levels of energetically less favourable signature peatland microbial processes such as fermentation, carbon fixation, methane metabolism, nitrogen fixation, assimilatory nitrate reduction and sulphate reduction. This provides a link of reduction in growth rates with a shift towards lower energy yielding pathways in healthier peatlands which likely leads to the lower organic matter decomposition and carbon accrual. Our DNA-based measurement of microbial growth only incorporates cell division but not other forms of growth. Specifically, under anoxic conditions, carbon taken up may not be entirely used for cell division or respiration, but a large proportion can be allocated to short-chain fatty acid production and release into the surrounding environment (Orsi et al., 2020). This may also explain the lower growth rates in the less oxic natural and restored peat communities. Microorganisms in the healthier sites also possessed lower capabilities for degradation of aromatic compounds and for recycling of microbial necromass. An approach similar to ours to link microbial traits to carbon cycling processes comparing a temperate bog and a fen highlighted similar findings in terms of metabolic pathways and substrate degradation capabilities (Richy et al., 2024). We also suggest that the growth rate reduction on restoration was larger in sites that were highly degraded by drainage or erosion, and it was larger when restoration age was greater. This needs to be tested further and can likely serve as a good indicator of microbial recovery following restoration.

We show that peat moisture content and oxygen availability which is dependent on the water table have a substantial influence on microbial ecophysiology and growth rates in the surface layers of peatlands along with other factors such as the vegetation and peat organic matter composition. Despite rewetting of peatlands having dramatic impacts on environmental conditions and microbial functions, restored peatlands in their surface layers continue to show the legacy of degradation even after ten years of restoration with likely lower resistance to reverse its carbon accrual functions when the water table is transiently lowered. The current strategy of restoring peatlands by raising the water table and re-vegetating seem to be effective in restoring microbial functions to the reference levels albeit with some differences. We provide a predictable association of microbial growth and metabolism to peatland ecosystem health at a landscape scale highlighting the key role of microbial processes in recovery of peatlands and their carbon accrual functions.

## Methods

### Field sites and sampling

We sampled seven peatland sites across mainland Britain (Figure 1a, Table S1) each with a degraded, restored, and near-natural treatment except Stean which lacked a near-natural area. The treatments were locally adjacent minimising the impact of underlying geology and climate. All sites were sampled between May and October 2021. The near-natural reference areas had no noticeable or known history of drainage features, ditches, or erosion. The length of time since drainage is unknown, but it is likely to have lasted for decades. Restoration was carried out on most sites for 9-12 years with two exceptions: restoration age was 6 years for Langwell and 3 years for Balmoral. Three replicates were sampled per site per treatment every 5 metres along a 10 metre transect. In drained and restored areas, samples were taken 2 m away from and parallel to drainage features. Samples were taken from similar peatland microtopography (lawns). Peat cores with a diameter of between 3-4 cm were taken using either a Russian peat corer or a standard soil corer excluding the top living vegetation. Cores were sliced into 5 cm sections, placed into sterile bags and cooled immediately. All equipment was washed with deionised water and cutting equipment with 70% ethanol between samples. Samples were frozen within 24 hours at -20 °C. Upon return to the lab, peat samples were thawed, and the top 10 cm sections were homogenised and aliquoted for different measurements; in some cases, aliquots were refrozen at -20 °C until analysis.

### Environmental characteristics

Oxygen concentration was measured using a fibre optic oxygen sensor (OXROB10, PyroScience, Germany) inserted into the peat core prior to homogenising. This was done separately for the 0-5 and 5-10 cm sections, and a mean of the two was taken to represent the topsoil dissolved oxygen concentration. While oxygen concentration estimates can be compromised due to the transport of samples to the lab, we observed clear patterns not just across the ecosystem health categories but also along the depth profile (data not shown) which suggest that the data are valid. Bulk density was measured as the volume displaced by frozen samples in a measuring cylinder (Chambers et al., 2010). Moisture content was measured gravimetrically expressed as a percentage of total weight. pH was measured on a slurry created using 25 ml deionised water added to 5 g of peat sample. Peat samples were dried and ball milled, and 10-12 mg of each sample were weighed into a tin capsule for elemental (C, N) analysis (NA 2500 Series, CE Instruments). Organic matter chemistry of peat soils was measured using Fourier Transform Infrared Spectroscopy (FTIR) with freeze dried and ball milled samples. Spectra were characterised between 4000 and 350 cm^-1^ through the averaging of 200 scans at 4 cm^-1^ using a Nicolet Magna-IR 550 FTIR spectrometer (Thermo Electron). Spectra were normalised by subtracting the minimum value and dividing by the average for each spectrum. To calculate the top cover of vegetation, a quadrat was placed over each sampling location and a photograph was taken. From the photograph, the area covered by main vegetation groups was calculated using ImageJ and results were expressed as percentages of total top cover for mosses, graminoids, heather, and other vegetation. Vegetation cover data was available for all but one site (Stean). The water table was measured at four sites (Langwell, Crocach, Moor House and Migneint) using Rugged TROLL 100 Water Level Loggers (In-Situ Inc.), which were installed inside metre long dipwell tubes in three locations per site per treatment. Loggers were installed between June 2021 and July 2022. The water table depth was measured for one year and expressed as the mean annual water table depth. This provides an indicative baseline water table for the sites and treatments (See https://github.com/NBellGroup/water-loggers/tree/main for full methods).

### Microbial growth rate measurement

Microbial community-level growth rates were determined by measuring ^18^O incorporation into total DNA (Spohn et al., 2016; Walker et al., 2018). An aliquot of 300 mg was weighed into 13 ml glass vials and preincubated at 15 °C for one week before the start of the experiment to allow microbial activity to stabilise indicated by soil respiration reaching basal rates. ^18^O-H_2_O (Elemtex 98% enrichment) was added to the samples to reach an enrichment of 30 at% of ^18^O in the final soil water. A natural abundance control was maintained with the same volume of MilliQ water. After 48 hours of incubation at 15 °C, the samples were harvested and frozen at -20°C. DNA was extracted from 250 mg of peat using the DNeasy PowerSoil Pro Kits (Qiagen) following manufacturer’s instructions. DNA extracts were dried in silver capsules overnight at 60°C. The ^18^O abundance and total oxygen content was measured using a High Temperature Conversion Analyser (TC/EA, Thermo Scientific) coupled to an isotope ratio mass spectrometer (Delta V Advantage, Thermo Scientific). Based on the amount of new DNA produced with ^18^O enrichment, we calculated the total new biomass production rate during the incubation period using a conversion factor (Spohn et al., 2016).

### Metagenomics sequencing and bioinformatics

DNA was extracted following the same protocol as described above. DNA quality and quantity was assessed using Nanodrop and Qubit fluorometric assays. Illumina libraries were prepared using NEBNext Ultra II FS DMA library prep kit according to manufacturer’s instructions. Sequencing was carried out on an Illumina Novaseq at the NERC Environmental Omics Facility. Adapter trimming and quality filtering (>Q20) of retrieved raw sequences was carried out using Cutapdapt version 1.2.1 and Sickle version 1.2; after trimming reads with a length of less than 15 bp were removed. The reads were assembled using metaSPAdes version 3.13.0 (Bankevich et al., 2012; Nurk et al., 2013); contigs of less than 1000 bp were removed. Prodigal was used to carry out gene-calling of metagenomic contigs (Hyatt et al., 2010). Contigs-based functional gene annotations were performed for CAZyme and KEGG genes using dbcan2 version 2.0.11 (Huang et al., 2018) and KOfamscan respectively version 1.2.0 (Aramaki et al., 2020). Functional gene abundances were normalised by total Prodigal-predicted protein-coding genes (i.e., total number of open reading frames) in metagenomic contigs to account for assembly bias and variation in sequencing depth.

### Genome binning

To achieve binning of high quality MAGs, we co-assembled by merging reads from three replicates from each site-treatment combination. Binning and quality control was carried out using MetaBAT2 version 2.15 (Kang et al., 2019). Binners were applied over the co-assemblies to extract bins. The binners were run using default parameters. CheckM version 1.2.2 was run over the collected bins or MAGs to determine completion, contamination, and bin abundances across the samples (Parks et al., 2015). Contamination is the percentage of a sequence in the MAG which belongs to a different species/strain and completeness of a MAG is estimated from the fraction of certain marker genes present within the genome. MAGs above 10% contamination and below 50% completeness were removed (Bowers et al., 2017). MAGs were dereplicated using dRep using default parameters except the use of skani for genome comparisons. Metagenome reads were then mapped to the MAGs using Bowtie2 v2.4.5 to obtain abundance and coverage statistics. SAMtools v1.17 was used to sort and index read mapping files. CoverM v0.6.1 was used to filter read mapping files to remove reads with <90% identity and to generate absolute mapped read counts.

### Statistical analyses and visualisations

All analyses were carried out in R v.4.3.1. Visualisations were created using ggplot2 v.3.4.3 and data manipulation through the tidyverse packages v.2.00 (Wickham, 2016; Wickham et al., 2019; Team, 2023). The map of sampling sites and peat extent was generated in R with the packages maps, mapdata and ggplot2 and land cover data for peat cover mapping was obtained from ArcGIS Hub at hub.arcgis.com/datasets/Defra::peaty-soils-location-england (England), hub.arcgis.com/datasets/theriverstrust::unified-peat-map-for-wales (Wales), and hub.arcgis.com/datasets/snh::carbon-and-peatland-2016-map (Scotland). Principal Component Analysis (PCA) on the environmental variables (moisture, oxygen concentration, moss cover, pH, carbon:nitrogen or C:N ratio and organic matter composition) was performed to examine how they contributed to system-level variation across the sites and treatments. Conductivity, vascular vegetation cover, and total carbon and nitrogen were excluded from this analysis because of collinearity with pH, moss over and C:N ratio, respectively. We calculated the percentage of variation which each variable explained across the first two principal components. PCA was performed using the PCA function from the package *FactoMineR* and plotted using the package factoextra (Kassambara and Mundt, 2020).

### Calculation of the ecosystem health index

An ecosystem health index was calculated as a weighted sum of standardized variables using the six key environmental variables that explained majority of the variation across all samples. These were dissolved oxygen concentration, C:N ratio, organic matter chemical composition as derived from the first PCA axis of FTIR spectroscopy which explained 50.72% of the variation across samples, moisture content, moss cover, and pH. Variables were first standardized (centred and scaled by a factor of 0.01 to reduce range) and scaled weights that were proportional to the sample variance explained by each factor were introduced. Some variables (oxygen, FTIR PC1 and C:N ratio) were reversed weighted to correct their direction of influence. Vegetation cover data was missing for one of the sites (Stean) for which the weights for the other variables were adjusted proportionally to maintain total weight.

### Site and treatment effect on environmental variables

We tested the effects of treatments (degraded, restored and natural) on environmental variables and microbial traits using the lmer function in lme4 package. To account for different underlying geology and climatic differences at each site we added site as a random variable in the model. A nested model structure of treatments nested within sites was also tested but its performance was similar to the model with site as a random variable. Estimated marginal means or least-squares means were used to study the effect of treatment, followed by pairwise analysis of the three treatments. Regression analysis of microbial growth rate against the ecosystem health index was performed using the lmer function with site as a random variable. A multiple linear regression model was fitted using the lm function to examine the relationship between microbial growth rate and the six key environmental factors: soil moisture, oxygen concentration, moss cover, pH, FTIR PC1 (representing organic matter composition), and C:N ratio. In addition, we also tested each factor individually using a simple linear regression model.

### Microbial community composition analysis

To examine the drivers of microbial community assembly we used Principal Coordinate Analysis (PCoA) based on Bray Curtis dissimilarity for the MAG abundance data across samples and treatments using the vegan package (OKSANEN, 2010). MAG abundance across sites and treatments was obtained as normalized trimmed mean genome coverages by mapping the metagenomic reads to MAG contigs, removing the top and bottom 5% of covered bases from coverage calculations, and dividing the resulting coverage values by sample sequencing depth. Regression of PCoA1 against the ecosystem health index was performed using the lmer function with site as a random variable.

### Microbial community-level functional analysis

We then tested to see whether key relevant functional pathways of peat microorganisms varied between treatments using community-level gene abundance obtained from contig-level information. We focused our analysis on genes for metabolic processes associated with five key pathways. We measured the sum gene abundance within the KEGG energy metabolism modules of nitrogen metabolism, sulphur metabolism, carbon fixation, and methane metabolism, and fermentation genes (Kanehisa et al., 2015). We measured the metabolic potential for organic matter degradation using CAZyme genes for specific substrates: cellulose, lignin, oligosaccharides, and microbial cell walls (Nuccio et al., 2020) and aromatic compound degradation genes from the KEGG module. We tested whether the abundance of these genes differed significantly across the treatments with the same mixed effect models used for the environmental variables and microbial growth rate. Regression analysis of gene abundances against the ecosystem health index was performed using the lmer function with site as a random variable.

### Functional analysis at the level of MAGs

To identify differentially abundant MAGs in each ecosystem health treatment, we ran DESeq2 (Love et al., 2014) using the absolute mapped read counts to MAGs for each sample site, separately. Differential abundance for each MAG was determined by likelihood ratio tests that compared model fits with and without ecosystem health treatment, using a maximum false-discovery rate adjusted *P* value of 0.05. To identify MAGs that were enriched in specific ecosystem health treatments, the differentially abundant MAGs from DESeq2 were hierarchically clustered according to their trimmed mean genome coverages, performed for each site separately. The filtered trimmed mean coverages were converted into abundances relative to the total abundance within each sample for clustering, and MAGs were assigned to clusters based on the Z-scores of their abundance compared to the means across all treatments. For the differentially abundant MAGs, we assessed metabolic functions using METABOLIC v4.0 with the METABOLIC-C.pl program. The “KEGGModuleHit” results were used to determine pathway presence/absence in MAGs. We focused on the different individual functional gene categories within nitrogen metabolism, sulphur metabolism, carbon fixation, methane metabolism, and fermentation.

## Data Availability

All raw sequencing data are publicly available in the NCBI Short Read Archive under BioProject accession PRJNA1203648. Whole assembled metagenomic contigs as well as high-quality prokaryotic metagenome-assembled genomes are available at the NCBI WGS using the same BioProject accession.

## Contributions

WP and AAM designed the research; AAM acquired funding, administered the project, and was the primary PhD supervisor of WP; WP, EK, and NB performed the soil sampling; WP performed most of the analytical work with the exception of the following analytical procedures; CM performed the FTIR analysis; EK performed the water table and vegetation cover analysis; AR facilitated the DNA ^18^O analysis for microbial growth measurements; WP, YW, JCK, and AAM performed the bioinformatics, statistical analyses, visualisations and data curation; RREA, RA, TCP, and AAM were involved in supervision of WP’s doctoral studies; WP and AAM drafted the manuscript; all authors were involved in critical revision, editing, and approval of the final version.

## Acknowledgments

We gratefully acknowledge the landowners, site managers, and researchers who enabled access to the field sites used in this study. We wish to thank Emily MacDonald, Nicole Cochrane and Hedda Weitz (University of Aberdeen), and Angela Main (James Hutton Institute) for technical support. We thank Antonio Riberio (University of Aberdeen) for bioinformatics support.

## Reference

Amacher, M.C., O’Neil, K.P., Perry, C.H., 2007. Soil vital signs: A new Soil Quality Index (SQI) for assessing forest soil health. U.S. Department of Agriculture, Forest Service, Rocky Mountain Research Station. doi:10.2737/rmrs-rp-65

Andersen, R., Chapman, S.J., Artz, R.R.E., 2012. Microbial communities in natural and disturbed peatlands: A review. doi:10.1016/j.soilbio.2012.10.003

Andersen, R., Farrell, C., Graf, M., Muller, F., Calvar, E., Frankard, P., Caporn, S., Anderson, P., 2017. An overview of the progress and challenges of peatland restoration in Western Europe. Restoration Ecology 25, 271–282. doi:10.1111/REC.12415

Andersen, R., Wells, C., Macrae, M., Price, J., 2013. Nutrient mineralisation and microbial functional diversity in a restored bog approach natural conditions 10 years post restoration. Soil Biology and Biochemistry 64, 37–47. doi:10.1016/j.soilbio.2013.04.004

Aramaki, T., Blanc-Mathieu, R., Endo, H., Ohkubo, K., Kanehisa, M., Goto, S., Ogata, H., 2020. KofamKOALA: KEGG Ortholog assignment based on profile HMM and adaptive score threshold. Bioinformatics 36, 2251–2252. doi:10.1093/bioinformatics/btz859

Artz, R.R.E., Chapman, S.J., Jean Robertson, A.H., Potts, J.M., Laggoun-Défarge, F., Gogo, S., Comont, L., Disnar, J.-R., Francez, A.-J., 2008. FTIR spectroscopy can be used as a screening tool for organic matter quality in regenerating cutover peatlands. Soil Biology and Biochemistry 40, 515–527. doi:10.1016/j.soilbio.2007.09.019

Averesch, N.J.H., Kracke, F., 2018. Metabolic Network Analysis of Microbial Methane Utilization for Biomass Formation and Upgrading to Bio-Fuels. Frontiers in Energy Research Volume 6-2018. doi:10.3389/fenrg.2018.00106

Bankevich, A., Nurk, S., Antipov, D., Gurevich, A.A., Dvorkin, M., Kulikov, A.S., Lesin, V.M., Nikolenko, S.I., Pham, S., Prjibelski, A.D., Pyshkin, A.V., Sirotkin, A.V., Vyahhi, N., Tesler, G., Alekseyev, M.A., Pevzner, P.A., 2012. SPAdes: A new genome assembly algorithm and its applications to single-cell sequencing. Journal of Computational Biology 19, 455–477. doi:10.1089/CMB.2012.0021/FORMAT/EPUB

Bell, S.L., Zimmerman, A.E., Stone, B.W., Chang, C.H., Blumer, M., Renslow, R.S., Propster, J.R., Hayer, M., Schwartz, E., Hungate, B.A., Hofmockel, K.S., 2023. Effects of warming on bacterial growth rates in a peat soil under ambient and elevated CO2. Soil Biology and Biochemistry 178, 108933. doi:10.1016/j.soilbio.2022.108933

Bengtsson, F., Rydin, H., Hájek, T., 2018. Biochemical determinants of litter quality in 15 species of Sphagnum. Plant and Soil 425, 161–176. doi:10.1007/S11104-018-3579-8/FIGURES/4

Bergman, I., Lundberg, P., Preston, C.M., Nilsson, M., 2000. Degradation of 13C–U–Glucose in Sphagnum majus Litter Responses to Redox, pH, and Temperature. Soil Science Society of America Journal 64, 1368–1381. doi:10.2136/sssaj2000.6441368x

Blüthgen, N., Dormann, C.F., Prati, D., Klaus, V.H., Kleinebecker, T., Hölzel, N., Alt, F., Boch, S., Gockel, S., Hemp, A., Müller, J., Nieschulze, J., Renner, S.C., Schöning, I., Schumacher, U., Socher, S.A., Wells, K., Birkhofer, K., Buscot, F., Oelmann, Y., Rothenwöhrer, C., Scherber, C., Tscharntke, T., Weiner, C.N., Fischer, M., Kalko, E.K.V., Linsenmair, K.E., Schulze, E.-D., Weisser, W.W., 2012. A quantitative index of land-use intensity in grasslands: Integrating mowing, grazing and fertilization. Basic and Applied Ecology 13, 207–220. doi:10.1016/j.baae.2012.04.001

Bowers, R.M., Kyrpides, N.C., Stepanauskas, R., Harmon-Smith, M., Doud, D., Reddy, T.B.K., Schulz, F., Jarett, J., Rivers, A.R., Eloe-Fadrosh, E.A., Tringe, S.G., Ivanova, N.N., Copeland, A., Clum, A., Becraft, E.D., Malmstrom, R.R., Birren, B., Podar, M., Bork, P., Weinstock, G.M., Garrity, G.M., Dodsworth, J.A., Yooseph, S., Sutton, G., Glöckner, F.O., Gilbert, J.A., Nelson, W.C., Hallam, S.J., Jungbluth, S.P., Ettema, T.J.G., Tighe, S., Konstantinidis, K.T., Liu, W.-T., Baker, B.J., Rattei, T., Eisen, J.A., Hedlund, B., McMahon, K.D., Fierer, N., Knight, R., Finn, R., Cochrane, G., Karsch-Mizrachi, I., Tyson, G.W., Rinke, C., Kyrpides, N.C., Schriml, L., Garrity, G.M., Hugenholtz, P., Sutton, G., Yilmaz, P., Meyer, F., Glöckner, F.O., Gilbert, J.A., Knight, R., Finn, R., Cochrane, G., Karsch-Mizrachi, I., Lapidus, A., Meyer, F., Yilmaz, P., Parks, D.H., Murat Eren, A., Schriml, L., Banfield, J.F., Hugenholtz, P., Woyke, T., The Genome Standards Consortium, 2017. Minimum information about a single amplified genome (MISAG) and a metagenome-assembled genome (MIMAG) of bacteria and archaea. Nature Biotechnology 35, 725–731. doi:10.1038/nbt.3893

Chambers, F.M., Beilman, D.W., Yu, Z., 2010. Methods for determining peat humification and for quantifying peat bulk density, organic matter and carbon content for palaeostudies of climate and peatland carbon dynamics 7, 1–10.

Elliott, D.R., Caporn, S.J.M., Nwaishi, F., Nilsson, R.H., Sen, R., 2015. Bacterial and Fungal Communities in a Degraded Ombrotrophic Peatland Undergoing Natural and Managed Re-Vegetation. PLOS ONE 10, e0124726. doi:10.1371/JOURNAL.PONE.0124726

Evans, C.D., Peacock, M., Baird, A.J., Artz, R.R.E., Burden, A., Callaghan, N., Chapman, P.J., Cooper, H.M., Coyle, M., Craig, E., Cumming, A., Dixon, S., Gauci, V., Grayson, R.P., Helfter, C., Heppell, C.M., Holden, J., Jones, D.L., Kaduk, J., Levy, P., Matthews, R., McNamara, N.P., Misselbrook, T., Oakley, S., Page, S.E., Rayment, M., Ridley, L.M., Stanley, K.M., Williamson, J.L., Worrall, F., Morrison, R., 2021. Overriding water table control on managed peatland greenhouse gas emissions. Nature 2021 593:7860 593, 548–552. doi:10.1038/s41586-021-03523-1

Geay, M.L., Lauga, B., Walcker, R., Jassey, V.E.J., 2024. A meta-analysis of peatland microbial diversity and function responses to climate change. Soil Biology and Biochemistry 189, 109287. doi:10.1016/J.SOILBIO.2023.109287

Gios, E., Verbruggen, E., Audet, J., Burns, R., Butterbach-Bahl, K., Espenberg, M., Fritz, C., Glatzel, S., Jurasinski, G., Larmola, T., Mander, Ü., Nielsen, C., Rodriguez, A.F., Scheer, C., Zak, D., Silvennoinen, H.M., 2024. Unraveling microbial processes involved in carbon and nitrogen cycling and greenhouse gas emissions in rewetted peatlands by molecular biology. Biogeochemistry 2024 167:4 167, 609–629. doi:10.1007/S10533-024-01122-6

Grinter, R., Greening, C., 2021. Cofactor F420: an expanded view of its distribution, biosynthesis and roles in bacteria and archaea. FEMS Microbiology Reviews 45, fuab021. doi:10.1093/femsre/fuab021

Gunnarsson, U., 2005. Global patterns of Sphagnum productivity. Journal of Bryology 27, 269–279. doi:10.1179/174328205X70029

Hamard, S., Planchenault, S., Walcker, R., Sytiuk, A., Le Geay, M., Küttim, M., Dorrepaal, E., Lamentowicz, M., Petchey, O.L., Robroek, B.J.M., Tuittila, E.-S., Barret, M., Céréghino, R., Delarue, F., Ferriol, J., Lafont Rapnouil, T., Leflaive, J., Le Roux, G., Jassey, V.E.J., 2025. Microbial photosynthesis mitigates carbon loss from northern peatlands under warming. Nature Climate Change 15, 436–443. doi:10.1038/s41558-025-02271-8

Holden, J., Chapman, P.J., Labadz, J.C., 2004. Artificial drainage of peatlands: hydrological and hydrochemical process and wetland restoration. http://Dx.Doi.Org/10.1191/0309133304pp403ra 28, p95–123. doi:10.1191/0309133304PP403RA

Huang, L., Zhang, H., Wu, P., Entwistle, S., Li, X., Yohe, T., Yi, H., Yang, Z., Yin, Y., 2018. dbCAN-seq: a database of carbohydrate-active enzyme (CAZyme) sequence and annotation. Nucleic Acids Research 46, D516. doi:10.1093/NAR/GKX894

Huang, Y., Ciais, P., Luo, Y., Zhu, D., Wang, Y., Qiu, C., Goll, D.S., Guenet, B., Makowski, D., De Graaf, I., Leifeld, J., Kwon, M.J., Hu, J., Qu, L., 2021. Tradeoff of CO2 and CH4 emissions from global peatlands under water-table drawdown. Nature Climate Change 11, 618–622. doi:10.1038/s41558-021-01059-w

Hyatt, D., Chen, G.-L., LoCascio, P.F., Land, M.L., Larimer, F.W., Hauser, L.J., 2010. Prodigal: prokaryotic gene recognition and translation initiation site identification. BMC Bioinformatics 11, 119. doi:10.1186/1471-2105-11-119

Joosten, H., Sirin, A., Couwenberg, J., Laine, J., Smith, P., 2016. The role of peatlands in climate regulation, in: Bonn, A., Allott, T., Evans, M., Joosten, H., Stoneman, R. (Eds.), Peatland Restoration and Ecosystem Services: Science, Policy and Practice, Ecological Reviews. Cambridge University Press, Cambridge, pp. 63–76. doi:10.1017/CBO9781139177788.005

Kalbitz, K., Geyer, S., 2002. Different effects of peat degradation on dissolved organic carbon and nitrogen. Organic Geochemistry 33, 319–326. doi:10.1016/S0146-6380(01)00163-2

Kanehisa, M., Sato, Y., Kawashima, M., Furumichi, M., Tanabe, M., 2015. KEGG as a reference resource for gene and protein annotation. Nucleic Acids Research 44, 457– 462. doi:10.1093/nar/gkv1070

Kang, D.D., Li, F., Kirton, E., Thomas, A., Egan, R., An, H., Wang, Z., 2019. MetaBAT 2: an adaptive binning algorithm for robust and efficient genome reconstruction from metagenome assemblies. PeerJ 7, e7359. doi:10.7717/peerj.7359

Kassambara, A., Mundt, F., 2020. factoextra: Extract and Visualize the Results of Multivariate Data Analyses.

Kitson, E., Bell, N.G.A., 2020. The Response of Microbial Communities to Peatland Drainage and Rewetting. A Review. Frontiers in Microbiology 11.

Klimkowska, A., Goldstein, K., Wyszomirski, T., Kozub, Ł., Wilk, M., Aggenbach, C., Bakker, J.P., Belting, H., Beltman, B., Blüml, V., Vries, Y.D., Geiger-Udod, B., Grootjans, A.P., Hedberg, P., Jager, H.J., Kerkhof, D., Kollmann, J., Pawlikowski, P., Pleyl, E., Reinink, W., Rydin, H., Schrautzer, J., Sliva, J., Stańko, R., Sundberg, S., Timmermann, T., Wołejko, L., Burg, R.F. van der, Hoek, D. van der, Diggelen, J.M.H. van, Heerden, A. van, Tweel, L. van, Vegelin, K., Kotowski, W., 2019. Are we restoring functional fens? – The outcomes of restoration projects in fens re-analysed with plant functional traits. PLOS ONE 14, e0215645. doi:10.1371/journal.pone.0215645

Köchy, M., Hiederer, R., Freibauer, A., 2015. Global distribution of soil organic carbon – Part 1: Masses and frequency distributions of SOC stocks for the tropics, permafrost regions, wetlands, and the world. SOIL 1, 351–365. doi:10.5194/soil-1-351-2015

Krüger, J.P., Leifeld, J., Glatzel, S., Szidat, S., Alewell, C., 2015. Biogeochemical indicators of peatland degradation – a case study of a temperate bog in northern Germany. Biogeosciences 12, 2861–2871. doi:10.5194/bg-12-2861-2015

Laiho, R., 2006. Decomposition in peatlands: Reconciling seemingly contrasting results on the impacts of lowered water levels. Soil Biology and Biochemistry 38, 2011–2024. doi:10.1016/J.SOILBIO.2006.02.017

Lehmann, J., Bossio, D.A., Kögel-Knabner, I., Rillig, M.C., 2020. The concept and future prospects of soil health. Nature Reviews Earth & Environment 1, 544–553. doi:10.1038/s43017-020-0080-8

Leifeld, J., Klein, K., Wüst-Galley, C., 2020. Soil organic matter stoichiometry as indicator for peatland degradation. Scientific Reports 10, 7634. doi:10.1038/s41598-020-64275-y

Leifeld, J., Menichetti, L., 2018. The underappreciated potential of peatlands in global climate change mitigation strategies. Nature Communications 2018 9:1 9, 1–7. doi:10.1038/s41467-018-03406-6

Leifeld, J., Wüst-Galley, C., Page, S., 2019. Intact and managed peatland soils as a source and sink of GHGs from 1850 to 2100. Nature Climate Change 2019 9:12 9, 945–947. doi:10.1038/s41558-019-0615-5

Li, Z., Lun, F., Liu, M., Xiao, X., Wang, C., Wang, L., Xu, Y., Qi, W., Sun, D., 2021. Rapid diagnosis of agricultural soil health: A novel soil health index based on natural soil productivity and human management. Journal of Environmental Management 277, 111402. doi:10.1016/j.jenvman.2020.111402

Lin, X., Tfaily, M.M., Green, S.J., Steinweg, J.M., Chanton, P., Imvittaya, A., Chanton, J.P., Cooper, W., Schadt, C., Kostka, J.E., 2014. Microbial Metabolic Potential for Carbon Degradation and Nutrient (Nitrogen and Phosphorus) Acquisition in an Ombrotrophic Peatland. doi:10.1128/AEM.00206-14

Lipson, D.A., Haggerty, J.M., Srinivas, A., Raab, T.K., Sathe, S., Dinsdale, E.A., 2013. Metagenomic Insights into Anaerobic Metabolism along an Arctic Peat Soil Profile. PLOS ONE 8, e64659. doi:10.1371/JOURNAL.PONE.0064659

Loisel, J., Gallego-Sala, A.V., Amesbury, M.J., Magnan, G., Anshari, G., Beilman, D.W., Benavides, J.C., Blewett, J., Camill, P., Charman, D.J., Chawchai, S., Hedgpeth, A., Kleinen, T., Korhola, A., Large, D., Mansilla, C.A., Mamp, J., Bellen, S., West, J.B., Yu, Z., Bubier, J.L., Garneau, M., Moore, T., Sannel, A.B.K., Page, S., Vamp, M., Bechtold, M., Brovkin, V., Cole, L.E.S., Chanton, J.P., Christensen, T.R., Davies, M.A., Vleeschouwer, F., Finkelstein, S.A., Frolking, S., Gaamp, M., Gandois, L., Girkin, N., Harris, L.I., Heinemeyer, A., Hoyt, A.M., Jones, M.C., Joos, F., Juutinen, S., Kaiser, K., Lacourse, T., Lamentowicz, M., Larmola, T., Leifeld, J., Lohila, A., Milner, A.M., Minkkinen, K., Moss, P., Naafs, B.D.A., Nichols, J., Oamp, J., Payne, R., Philben, M., Piilo, S., Quillet, A., Ratnayake, A.S., Roland, T.P., Sjamp, S., Sonnentag, O., Swindles, G.T., Swinnen, W., Talbot, J., Treat, C., Valach, A.C., Wu, J., 2021. Expert assessment of future vulnerability of the global peatland carbon sink. Nature Climate Change. doi:10.1038/s41558-020-00944-0

Love, M.I., Huber, W., Anders, S., 2014. Moderated estimation of fold change and dispersion for RNA-seq data with DESeq2. Genome Biology 15, 550. doi:10.1186/s13059-014-0550-8

Martin, W.F., 2020. Older Than Genes: The Acetyl CoA Pathway and Origins. Frontiers in Microbiology Volume 11-2020.

McCarter, C.P.R., Rezanezhad, F., Quinton, W.L., Gharedaghloo, B., Lennartz, B., Price, J., Connon, R., Van Cappellen, P., 2020. Pore-scale controls on hydrological and geochemical processes in peat: Implications on interacting processes. Earth-Science Reviews 207, 103227. doi:10.1016/j.earscirev.2020.103227

Negassa, W., Eckhardt, K.-U., Regier, T., Leinweber, P., 2021. Dissolved organic matter concentration, molecular composition, and functional groups in contrasting management practices of peatlands. Journal of Environmental Quality 50, 1364–1380. doi:10.1002/jeq2.20284

Nuccio, E.E., Starr, E. Karaoz, • Ulas, Eoin, •, Brodie, L., Zhou, • Jizhong, Tringe S.G., Malmstrom, R.R., Woyke, T., Banfield, J.F., Firestone, M.K., Pett-Ridge, J., 2020. Niche differentiation is spatially and temporally regulated in the rhizosphere. The ISME Journal 14, 999–1014. doi:10.1038/s41396-019-0582-x

Nurk, S., Bankevich, A., Antipov, D., Gurevich, A., Korobeynikov, A., Lapidus, A., Prjibelsky, A., Pyshkin, A., Sirotkin, A., Sirotkin, Y., Stepanauskas, R., McLean, J., Lasken, R., Clingenpeel, S.R., Woyke, T., Tesler, G., Alekseyev, M.A., Pevzner, P.A., 2013. Assembling Genomes and Mini-metagenomes from Highly Chimeric Reads. Lecture Notes in Computer Science (Including Subseries Lecture Notes in Artificial Intelligence and Lecture Notes in Bioinformatics) 7821 LNBI, 158–170. doi:10.1007/978-3-642-37195-0_13

Oksanen, J., 2010. Vegan : community ecology package. http://Vegan.r-Forge.r-Project.Org/.

Orsi, W.D., Schink, B., Buckel, W., Martin, W.F., 2020. Physiological limits to life in anoxic subseafloor sediment. FEMS Microbiology Reviews 44, 219–231. doi:10.1093/femsre/fuaa004

Parks, D.H., Imelfort, M., Skennerton, C.T., Hugenholtz, P., Tyson, G.W., 2015. CheckM: assessing the quality of microbial genomes recovered from isolates, single cells, and metagenomes. Genome Research 25, 1043–1055. doi:10.1101/gr.186072.114

Pester, M., Knorr, K.-H., Friedrich, M.W., Wagner, M., Loy, A., 2012. Sulfate-reducing microorganisms in wetlands – fameless actors in carbon cycling and climate change. Frontiers in Microbiology Volume 3-2012.

Potvin, L.R., Kane, E.S., Chimner, R.A., Kolka, R.K., Lilleskov, E.A., 2015. Effects of water table position and plant functional group on plant community, aboveground production, and peat properties in a peatland mesocosm experiment (PEATcosm). Plant and Soil 387, 277–294. doi:10.1007/s11104-014-2301-8

Poulter, B., Fluet-Chouinard, E., Hugelius, G., Koven, C., Fatoyinbo, L., Page, S.E., Rosentreter, J.A., Smart, L.S., Taillie, P.J., Thomas, N., Zhang, Z., Wijedasa, L.S., 2021. A Review of Global Wetland Carbon Stocks and Management Challenges, in: Wetland Carbon and Environmental Management, Geophysical Monograph Series. pp. 1–20. doi:10.1002/9781119639305.ch1

Rajakaruna, S., Makke, G., Grachet, N.G., Ayala-Ortiz, C., Bouranis, J., Hoyt, D.W., Toyoda, J., Denis, E.H., Moran, J.J., Song, T., Sun, X., Eder, E.K., Wong, A.R., Chu, R., Heyman, H., Kolton, M., Chanton, J.P., Wilson, R.M., Kostka, J., Tfaily, M.M., 2024. Adding labile carbon to peatland soils triggers deep carbon breakdown. Communications Earth & Environment 5, 792. doi:10.1038/s43247-024-01954-y

Richy, E., Cabello-Yeves, P.J., Hernandes-Coutinho, F., Rodriguez-Valera, F., González-Álvarez, I., Gandois, L., Rigal, F., Lauga, B., 2024. How microbial communities shape peatland carbon dynamics: New insights and implications. Soil Biology and Biochemistry 191, 109345. doi:10.1016/j.soilbio.2024.109345

Ritson, J.P., Alderson, D.M., Robinson, C.H., Burkitt, A.E., Heinemeyer, A., Stimson, A.G., Gallego-Sala, A., Harris, A., Quillet, A., Malik, A.A., 2021. Towards a microbial process-based understanding of the resilience of peatland ecosystem service provisioning–a research agenda. Science of the Total Environment 759, 143467.

Rowland, J.A., Bracey, C., Moore, J.L., Cook, C.N., Bragge, P., Walsh, J.C., 2021. Effectiveness of conservation interventions globally for degraded peatlands in cool-climate regions. Biological Conservation 263, 109327. doi:10.1016/j.biocon.2021.109327

Schimel, J., Schaeffer, S.M., 2012. Microbial control over carbon cycling in soil. Frontiers in Microbiology 3.

Silins, U., Rothwell, R.L., 2011. Spatial patterns of aerobic limit depth and oxygen diffusion rate at two peatlands drained for forestry in Alberta. 10.1139/X98-179 29, 53–61. doi:10.1139/X98-179

Sokol, N.W., Slessarev, E., Marschmann, G.L., Nicolas, A., Blazewicz, S.J., Brodie, E.L., Firestone, M.K., Foley, M.M., Hestrin, R., Hungate, B.A., Koch, B.J., Stone, B.W., Sullivan, M.B., Zablocki, O., Trubl, G., McFarlane, K., Stuart, R., Nuccio, E., Weber, P., Jiao, Y., Zavarin, M., Kimbrel, J., Morrison, K., Adhikari, D., Bhattacharaya, A., Nico, P., Tang, J., Didonato, N., Paša-Tolić, L., Greenlon, A., Sieradzki, E.T., Dijkstra, P., Schwartz, E., Sachdeva, R., Banfield, J., Pett-Ridge, J., LLNL Soil Microbiome Consortium, 2022. Life and death in the soil microbiome: how ecological processes influence biogeochemistry. Nature Reviews Microbiology 20, 415–430. doi:10.1038/s41579-022-00695-z

Spohn, M., Klaus, K., Wanek, W., Richter, A., 2016. Microbial carbon use efficiency and biomass turnover times depending on soil depth – Implications for carbon cycling. Soil Biology and Biochemistry 96, 74–81. doi:10.1016/j.soilbio.2016.01.016

Strack, M., Waddington, J.M., 2007. Response of peatland carbon dioxide and methane fluxes to a water table drawdown experiment. Global Biogeochemical Cycles 21. doi:10.1029/2006GB002715

Swindles, G.T., Morris, P.J., Mullan, D.J., Payne, R.J., Roland, T.P., Amesbury, M.J., Lamentowicz, M., Turner, T.E., Gallego-Sala, A., Sim, T., Barr, I.D., Blaauw, M., Blundell, A., Chambers, F.M., Charman, D.J., Feurdean, A., Galloway, J.M., Gałka, M., Green, S.M., Kajukało, K., Karofeld, E., Korhola, A., Lamentowicz, Ł., Langdon, P., Marcisz, K., Mauquoy, D., Mazei, Y.A., McKeown, M.M., Mitchell, E.A.D., Novenko, E., Plunkett, G., Roe, H.M., Schoning, K., Sillasoo, Ü., Tsyganov, A.N., van der Linden, M., Väliranta, M., Warner, B., 2019. Widespread drying of European peatlands in recent centuries. Nature Geoscience 12, 922–928. doi:10.1038/s41561-019-0462-z

Sytiuk, A., Céréghino, R., Hamard, S., Delarue, F., Guittet, A., Barel, J.M., Dorrepaal, E., Küttim, M., Lamentowicz, M., Pourrut, B., Robroek, B.J.M., Tuittila, E.-S., Jassey, V.E.J., 2022. Predicting the structure and functions of peatland microbial communities from Sphagnum phylogeny, anatomical and morphological traits and metabolites. Journal of Ecology 110, 80–96. doi:10.1111/1365-2745.13728

Team, R.C., 2023. R: A Language and Environment for Statistical Computing.

Tiemeyer, B., Borraz, E.A., Augustin, J., Bechtold, M., Beetz, S., Beyer, C., Drösler, M., Ebli, M., Eickenscheidt, T., Fiedler, S., Förster, C., Freibauer, A., Giebels, M., Glatzel, S., Heinichen, J., Hoffmann, M., Höper, H., Jurasinski, G., Leiber-Sauheitl, K., Peichl-Brak, M., Roßkopf, N., Sommer, M., Zeitz, J., 2016. High emissions of greenhouse gases from grasslands on peat and other organic soils. Global Change Biology 22, 4134–4149. doi:10.1111/GCB.13303

Turetsky, M.R., Benscoter, B., Page, S., Rein, G., van der Werf, G.R., Watts, A., 2015. Global vulnerability of peatlands to fire and carbon loss. Nature Geoscience 8, 11–14. doi:10.1038/ngeo2325

Turetsky, M.R., Donahue, W.F., Benscoter, B.W., 2011. Experimental drying intensifies burning and carbon losses in a northern peatland. Nature Communications 2, 514. doi:10.1038/ncomms1523

Urbanová, Z., Bárta, J., 2016. Effects of long-term drainage on microbial community composition vary between peatland types. Soil Biology and Biochemistry 92, 16–26. doi:10.1016/J.SOILBIO.2015.09.017

Waddington, J.M., Morris, P.J., Kettridge, N., Granath, G., Thompson, D.K., Moore, P.A., 2015. Hydrological feedbacks in northern peatlands. Ecohydrology 8, 113–127. doi:10.1002/ECO.1493

Walker, T.W.N., Kaiser, C., Strasser, F., Herbold, C.W., Leblans, N.I.W., Woebken, D., Janssens, I.A., Sigurdsson, B.D., Richter, A., 2018. Microbial temperature sensitivity and biomass change explain soil carbon loss with warming. Nature Climate Change 8, 885–889. doi:10.1038/s41558-018-0259-x

Wang, H., Tian, J., Chen, H., Ho, M., Vilgalys, R., Bu, Z.-J., Liu, X., Richardson, C.J., 2021. Vegetation and microbes interact to preserve carbon in many wooded peatlands. Communications Earth & Environment 2, 1–8. doi:10.1038/s43247-021-00136-4

Wang, X.-N., Sun, G.-X., Zhu, Y.-G., 2017. Thermodynamic energy of anaerobic microbial redox reactions couples elemental biogeochemical cycles. Journal of Soils and Sediments 17, 2831–2846. doi:10.1007/s11368-017-1767-4

Wickham, H., 2016. ggplot2: elegant graphics for data analysis.

Wickham, H., Averick, M., Bryan, J., Chang, W., François, R., Grolemund, G., Hayes, A., Henry, L., Hester, J., Kuhn, M., Pedersen, T.L., Miller, E., Bache, S.M., Müller, K., Ooms, J., Robinson, D., Seidel, D.P., 2019. Welcome to the tidyverse.

Wilhelm, R.C., Amsili, J.P., Kurtz, K.S.M., van Es, H.M., Buckley, D.H., 2023. Ecological insights into soil health according to the genomic traits and environment-wide associations of bacteria in agricultural soils. ISME Communications 3, 1. doi:10.1038/s43705-022-00209-1

Wilkinson, S.L., Andersen, R., Moore, P.A., Davidson, S.J., Granath, G., Waddington, J.M., 2023. Wildfire and degradation accelerate northern peatland carbon release. Nature Climate Change 13, 456–461. doi:10.1038/s41558-023-01657-w

Worrall, F., Gibson, H.S., Hopkins, J., Young, J., Lyndsay, D., Lopez-Soldana, G., 2025. Using earth observation to develop a health index for peatlands. Science of The Total Environment 970, 178956. doi:10.1016/j.scitotenv.2025.178956

Xu, J., Morris, P.J., Liu, J., Holden, J., 2018. PEATMAP: Refining estimates of global peatland distribution based on a meta-analysis. CATENA 160, 134–140. doi:10.1016/j.catena.2017.09.010

Xue, W., Ma, H., Xiang, M., Tian, J., Liu, X., 2023. From Sphagnum to shrub: Increased acidity reduces peat bacterial diversity and keystone microbial taxa imply peatland degradation. Land Degradation & Development 34, 5259–5272. doi:10.1002/ldr.4842

Yang, T., Jiang, J., He, Q., Shi, F., Jiang, H., Wu, H., He, C., 2025. Impact of drainage on peatland soil environments and greenhouse gas emissions in Northeast China. Scientific Reports 15, 8320. doi:10.1038/s41598-025-92655-9

Yin, T., Feng, M., Qiu, C., Peng, S., 2022. Biological Nitrogen Fixation and Nitrogen Accumulation in Peatlands. Frontiers in Earth Science 10, 670867. doi:10.3389/FEART.2022.670867/BIBTEX

Yu, Z., Loisel, J., Brosseau, D.P., Beilman, D.W., Hunt, S.J., 2010. Global peatland dynamics since the Last Glacial Maximum. Geophysical Research Letters 37. doi:10.1029/2010GL043584

Zeh, L., Igel, M.T., Schellekens, J., Limpens, J., Bragazza, L., Kalbitz, K., 2020. Vascular plants affect properties and decomposition of moss-dominated peat, particularly at elevated temperatures. Biogeosciences 17, 4797–4813. doi:10.5194/bg-17-4797-2020

Zhang, Z., Furman, A., 2021. Soil redox dynamics under dynamic hydrologic regimes - A review. Science of The Total Environment 763, 143026. doi:10.1016/j.scitotenv.2020.143026

Zhong, Y., Jiang, M., Middleton, B.A., 2020. Effects of water level alteration on carbon cycling in peatlands. Ecosystem Health and Sustainability 6. doi:10.1080/20964129.2020.1806113

