## Supplementary figures and tables for "Recovery of microbial ecophysiology and carbon accrual functions in peatlands under restoration"

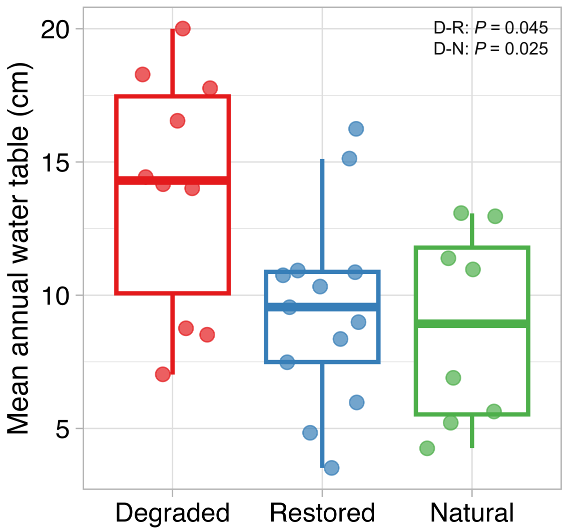


Figure S1: Mean annual water table for a period between June 2021 and July 2022 across the three treatments measured at four sites (Langwell, Crocach, Moor House and Migneint). Only significant pairwise differences in water table across the three treatments are highlighted with *P* values calculated using mixed effects models with site as a random effect (D: degraded, R: restored, and N: natural). Thick horizontal lines in the boxplots show median, boxes show interquartile ranges and vertical whiskers extend 1.5 times the inter quartile range.

**Table S1:** Study sites with their location, degradation status, and climatic data. Drained refers to the lowering of the water table through the digging of ditches which lead to the runoff of water. Eroded refers to the lowering of the water table through naturally formed ditches that arose due to other man-made disturbances such as fire or loss of vegetation. Mean annual precipitation (MAP) and mean annual temperature (MAT) were obtained from the Met office HadUK-Grid (Hollis et al., 2019).

| Site | Site data | Latitude Longitude | Degradation status |
|---|---|---|---|
| **Migneint** | MAP: 2181 mm | 52.99565 N 3.81652 W | Drained in the 1970's |
| Sampled: | MAT: 8 °C | 52.99835 N 3.80032 W | Restored 2011: through ditch blocking |
| May 2021 | Elevation: 453 m | 52.96932 N 3.81616 W | Near-natural |
| **Crocach** | MAP: 1258 mm | 58.39304 N 4.00182 W | Drained 1960s-70s |
| Sampled: | MAT: 7.1 °C | 58.38789 N 3.99658 W | Restored 2009 through ditch blocking |
| June 2021 | Elevation: 189 m | 58.39304 N 4.00182 W | Near-natural |
| **Stean** | MAP: 1229 mm | 54.1308 N 1.95015 W | Drained and heavily eroded 1960's-90's |
| Sampled: | MAT: 8 °C | 54.13559 N 1.92875 W | Restored 2012 through ditch blocking |
| June 2021 | Elevation: 530 m | | Near-natural not available |
| **Bowness** | MAP: 953 mm | 54.93297 N 3.23945 W | Drained 1970's |
| Sampled: | MAT: 9.6 °C | 54.92725 N 3.24204 W | Restored 2012 through ditch blocking |
| September 2021 | Elevation: 66 m | 54.93046 N 3.24013 W | Near-natural |
| **Moors House** | MAP: 1699 mm | 54.69166 N 2.38228 W | Eroded and drained 1960s |
| Sampled: | MAT: 8 °C | 54.69166 N 2.38124 W | Restored 2009 through ditch blocking |
| September 2021 | Elevation: 571 m | 54.69457 N 2.37661 W | Near-natural |
| **Langwell** | MAP: 1223 mm | 58.19629 N 3.61489 W | Drained 1970's |
| Sampled: | MAT: 7 °C | 58.20588 N 3.5552 W | Restored 2016 through ditch blocking |
| June 2021 | Elevation: 255 m | 58.2622 N 3.67514 W | Near-natural |
| **Balmoral** | MAP: 1412 mm | 56.92341 N 3.15831 W | Eroded since ~1950's |
| Sampled: | MAT: 5.5 °C | 56.92414 N 3.15579 W | Restored 2018 reprofiling |
| October 2021 | Elevation: 695 m | 56.92341 N 3.67514 W | Near-natural |

Reference:

Hollis D, McCarthy MP, Kendon M, Legg T, Simpson I. HadUK-Grid—A new UK dataset of gridded climate observations. *Geosci Data J.* 2019; 6: 151-159.

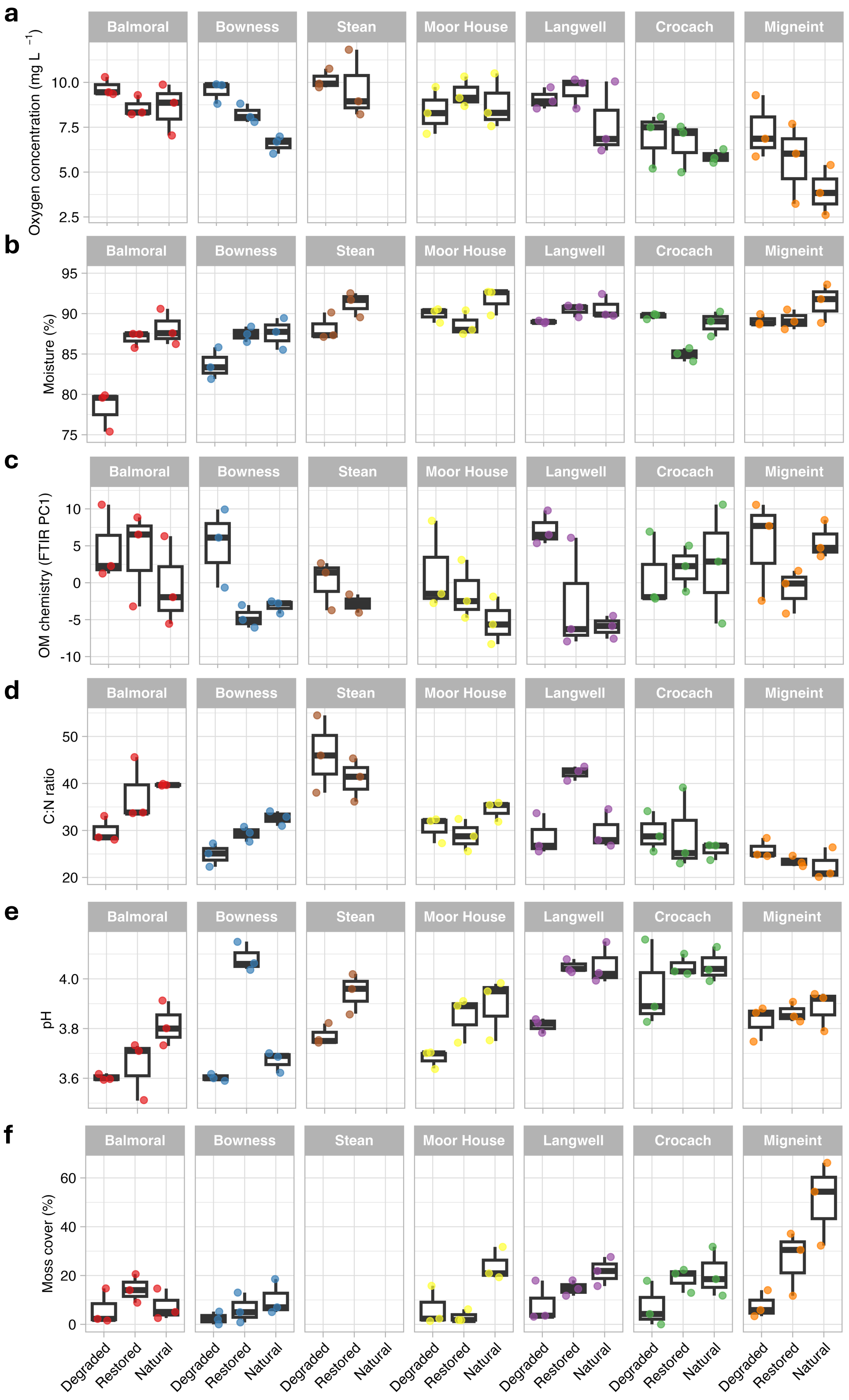


Figure S2: Environmental variables for all samples shown across individual sites and treatments of degraded, restored and natural peatlands for a) oxygen concentration, b) moisture content, c) organic matter chemical composition represented by FTIR PC1, d) C:N ratio of the organic matter, e) pH, and f) proportion of moss cover in the aboveground vegetation. Thick horizontal lines in the boxplots show median, boxes show interquartile ranges and vertical whiskers extend 1.5 times the inter quartile range.

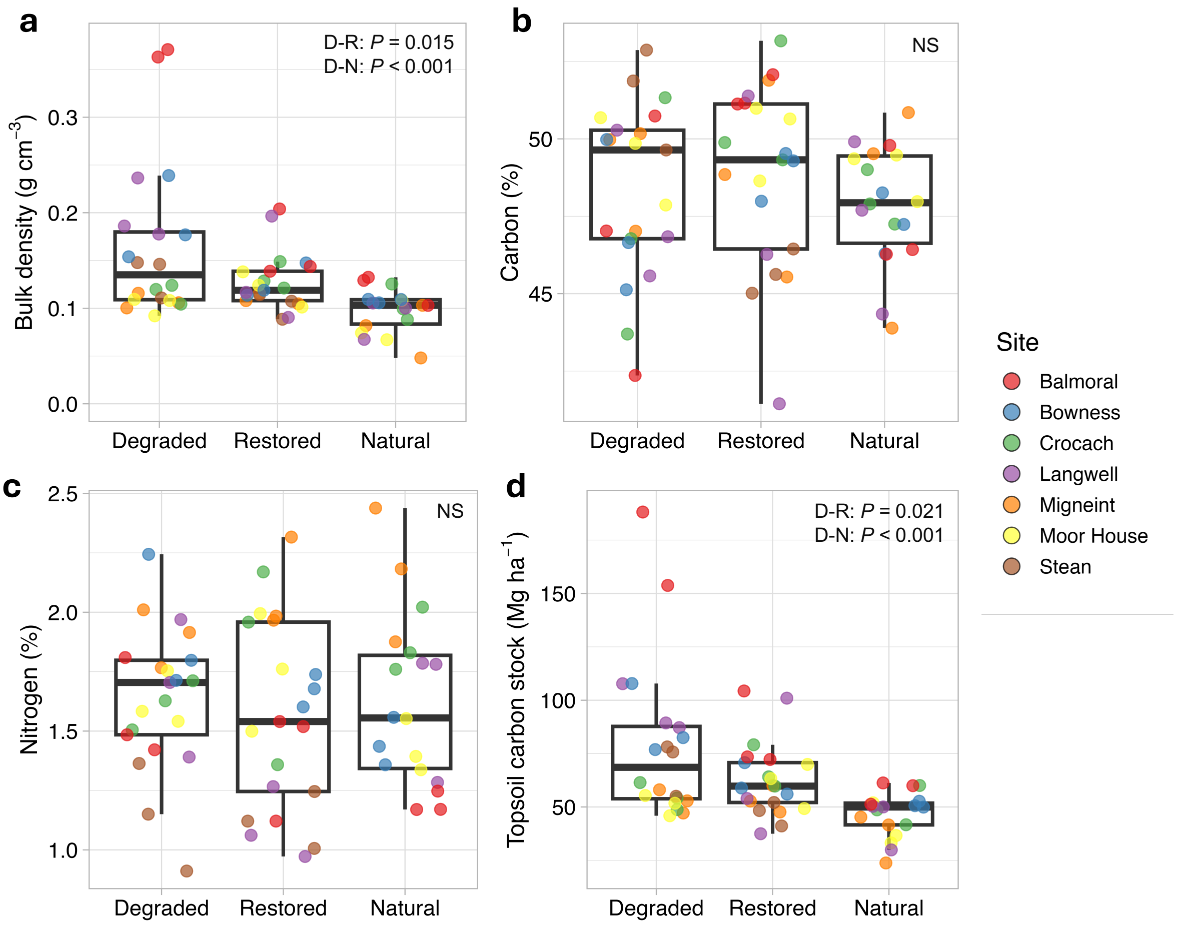


Figure S3: Environmental variables for all samples shown across the treatments of degraded, restored and natural peatlands for a) bulk density, b) carbon concentration, c) nitrogen concentration, and d) topsoil carbon stock. Only significant pairwise differences in environmental variables across the three treatments are highlighted with *P* values calculated using mixed effects models with site as a random effect (D: degraded, R: restored, N: natural, NS: not significant). Thick horizontal lines in the boxplots show median, boxes show interquartile ranges and vertical whiskers extend 1.5 times the inter quartile range.

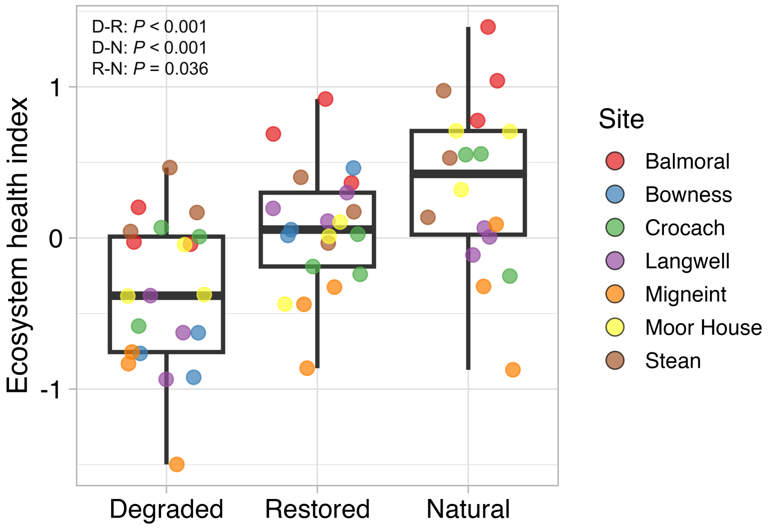


Figure S4: The ecosystem health index for all samples shown across the treatments of degraded, restored and natural peatlands. Only significant pairwise differences in the index across the three treatments are highlighted with *P* values calculated using mixed effects models with site as a random effect (D: degraded, R: restored, and N: natural). Thick horizontal lines in the boxplots show median, boxes show interquartile ranges and vertical whiskers extend 1.5 times the inter quartile range.

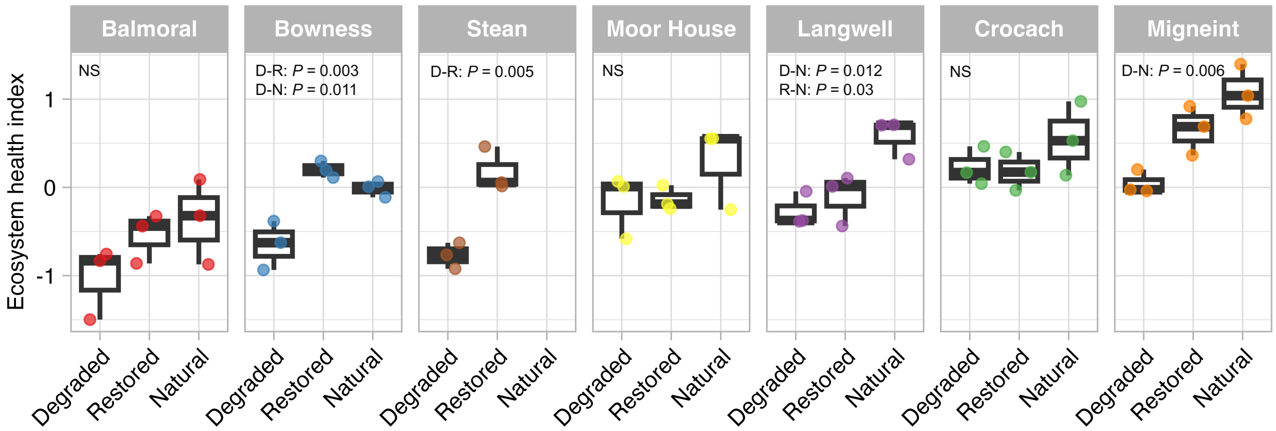


Figure S5: The ecosystem health index shown across individual sites and the treatments of degraded, restored and natural peatlands. Only significant pairwise differences in the index across the three treatments at individual sites are highlighted with *P* values calculated using mixed effects models (D: degraded, R: restored, N: natural, and NS: not significant). Thick horizontal lines in the boxplots show median, boxes show interquartile ranges and vertical whiskers extend 1.5 times the inter quartile range.

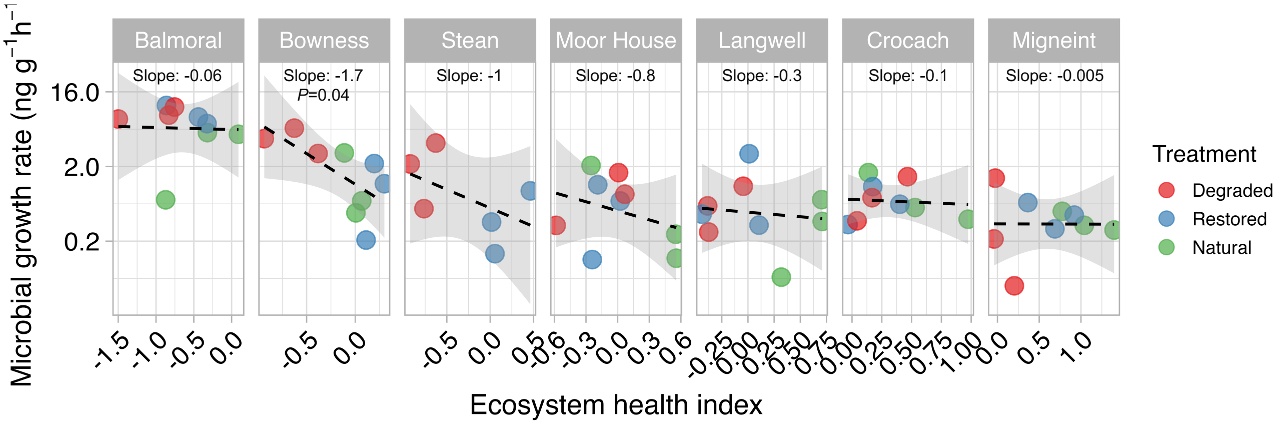


Figure S6: Microbial growth rates regressed against ecosystem heath index for all samples plotted for each site. The dashed line fits a linear regression model to the data with the shaded area showing a 95% confidence interval; the slope and *P* value were derived from a linear mixed-effects model. *P* values are only shown when significant (*P*<0.05).

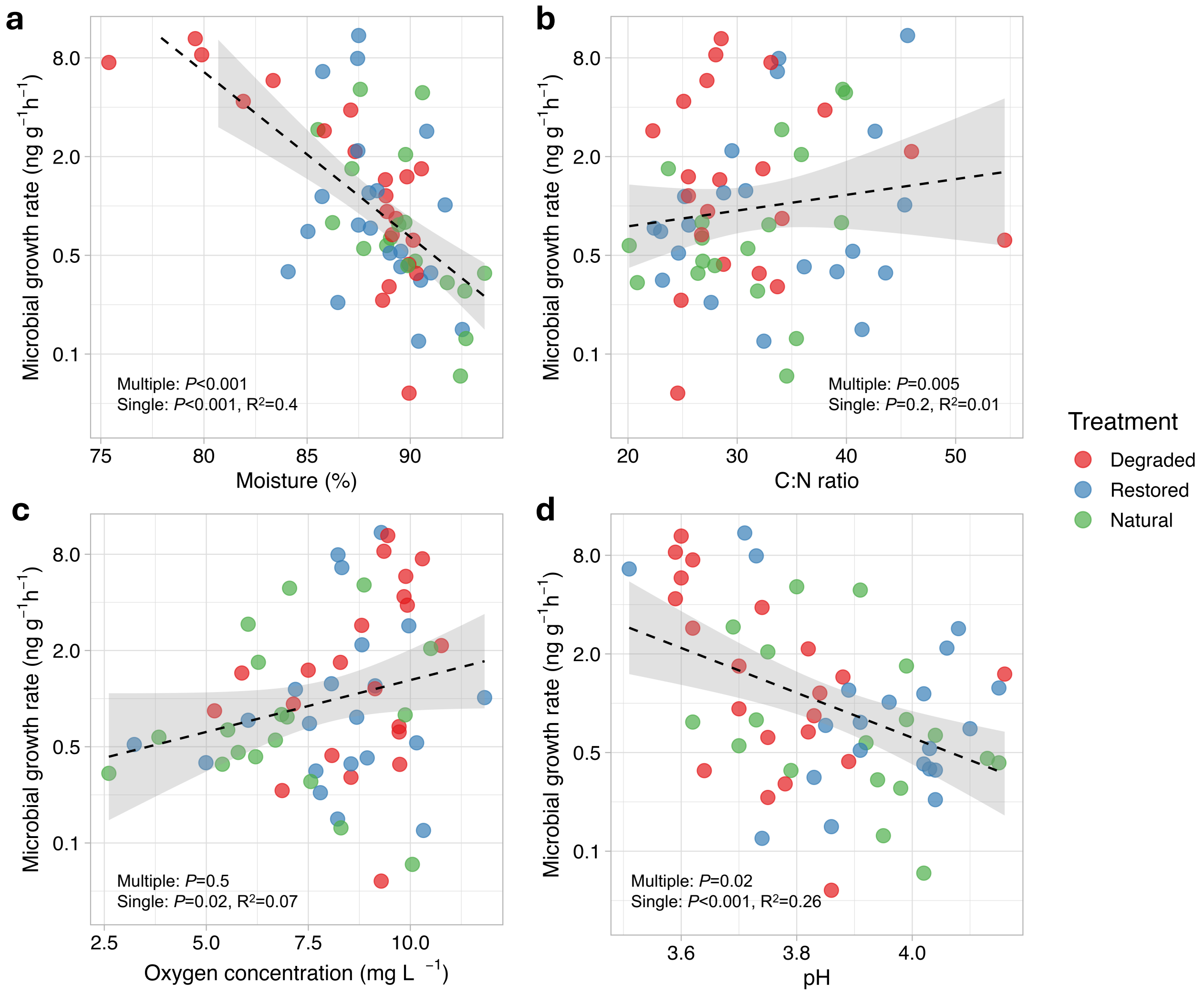


Figure S7: Microbial growth rates regressed against a) moisture content, b) peat organic matter C:N ratio, c) oxygen concentration, and d) peat pH. The dashed line fits a linear regression model to the data with the shaded area showing a 95% confidence interval. *P* values are displayed from a combined multiple linear regression model representing the variation in growth rate due to the six key environmental factors. *P* and $R^2$ values are also displayed from linear regression models of growth rate against the six environmental factors individually. Regression of growth rate with FTIR PC1 and moss cover were not significant (*P*>0.05) in any models.

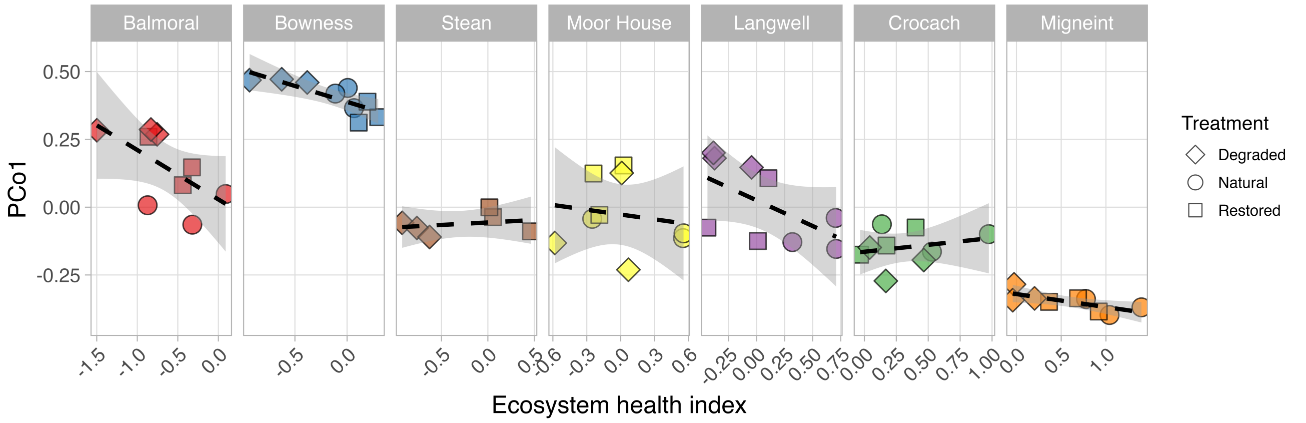


Figure S8: PCo1 from the PCoA analysis of microbial community composition regressed against ecosystem heath index for all samples plotted for each site. The dashed line fits a linear regression model to the data with the shaded area showing a 95% confidence interval. PCoA analysis was Bray Curtis distance-based for microbial community composition based on variation of MAG abundance across all samples.

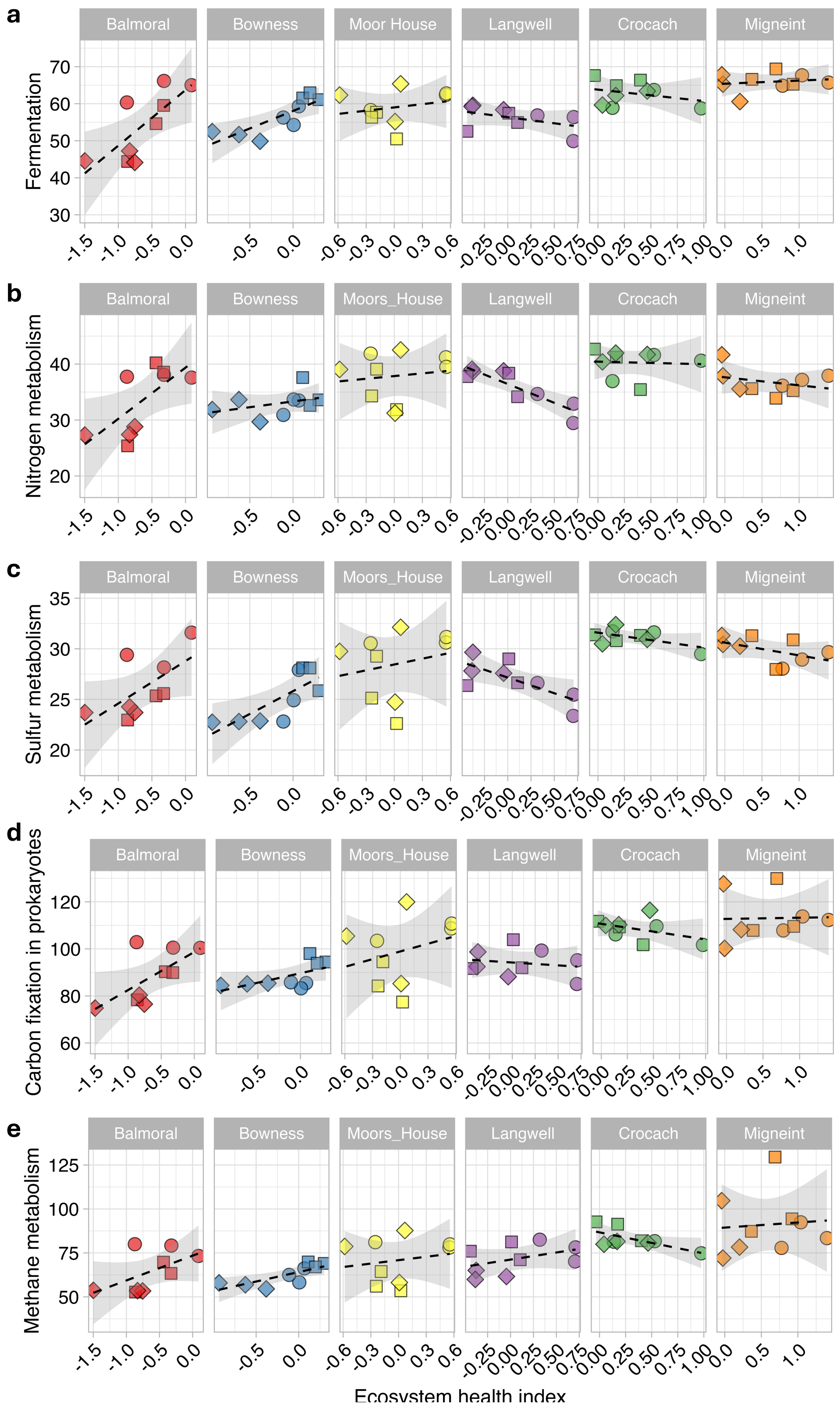


Figure S9: Gene abundances (as normalised reads ×1000) regressed against ecosystem heath index plotted for each site for key functional categories of metabolic pathways: a) fermentation, b) nitrogen metabolism, c) sulphur metabolism, d) carbon fixation in prokaryotes, and e) methane metabolism. The dashed line fits a linear regression model to the data with the shaded area showing a 95% confidence interval. The Stean site was excluded from this analysis. Point shapes represent treatments as per figure S8.

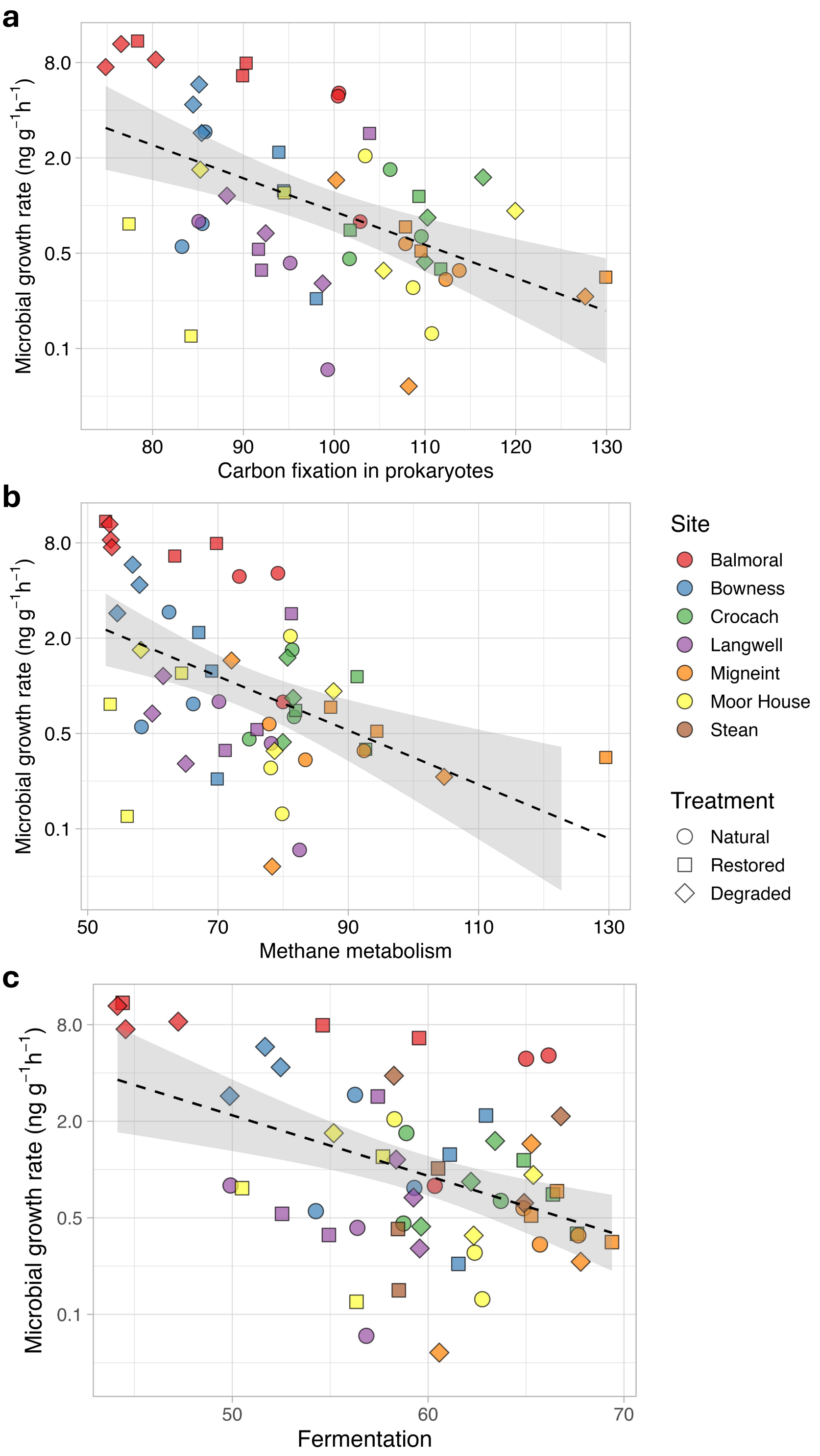


Figure S10: Microbial growth rates regressed against gene abundances for the metabolic pathways of significance in our sites a) carbon fixation in prokaryotes, b) methane metabolism, and c) fermentation. The dashed line fits a linear regression model to the data with the shaded area showing a 95% confidence interval.

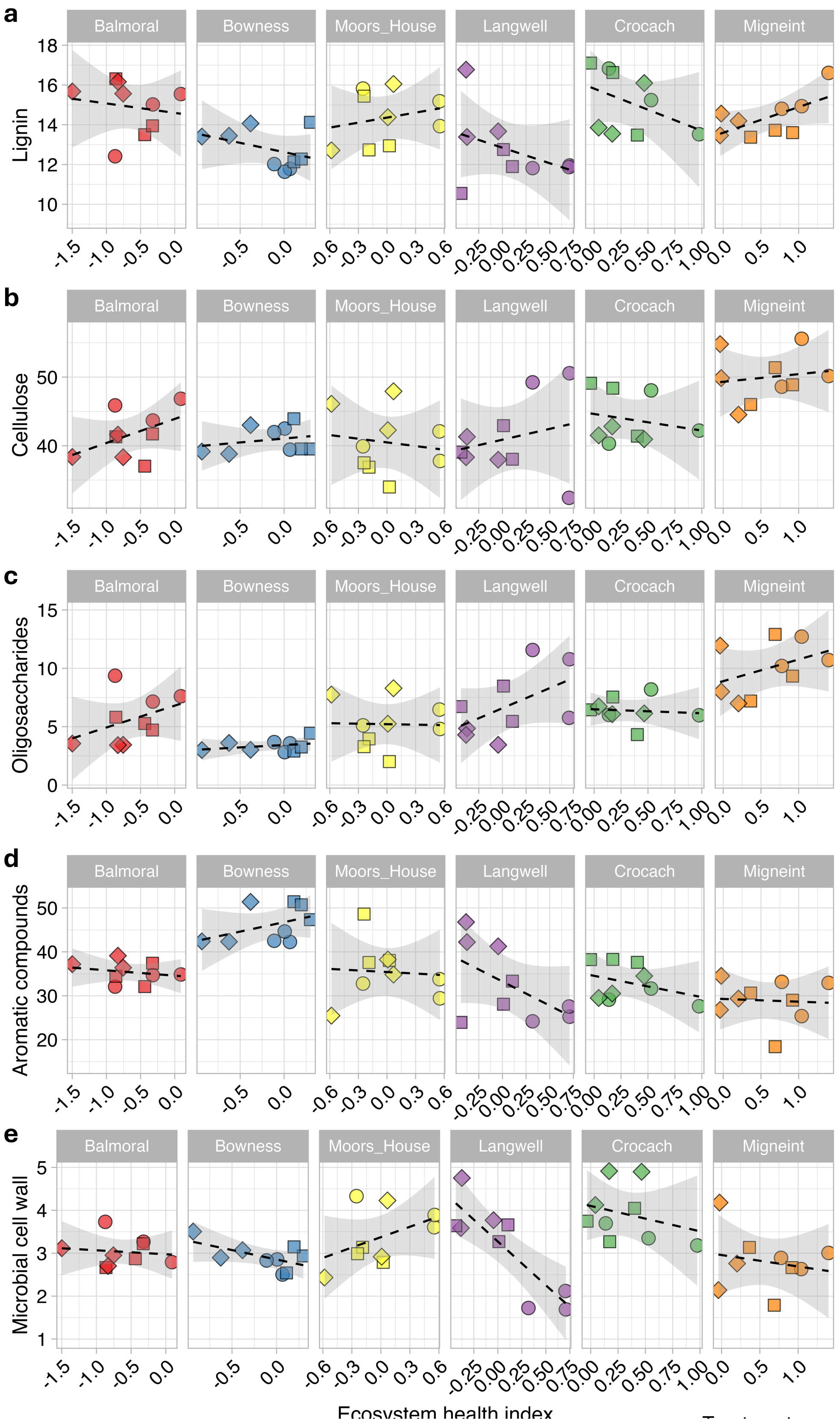


Figure S11: Regression against ecosystem heath index of abundances (as normalised reads ×1000) of genes for decomposition of a) lignin, b) cellulose, c) oligosaccharides, d) aromatic compounds, and e) microbial cell wall. The dashed line fits a linear regression model to the data with the shaded area showing a 95% confidence interval. The Stean site was excluded from this analysis. Point shapes represent treatments as per figure S8.
